# Combining FRET and super-resolution microscopy reveals kinase activation and mitochondrial activity at the nanoscale

**DOI:** 10.1101/2025.10.10.681576

**Authors:** Nicolas Y. Jolivet, Pierre-Jean Desmaison, Xavier Pinson, Arthur Masson, Olivier Delalande, Giulia Bertolin

## Abstract

Protein kinases are key regulators of intracellular signaling, showing activities in specific subcellular compartments and in micro- or nano-domains. Genetically encoded biosensors based on Förster’s Resonance Energy Transfer (FRET) are powerful tools to track kinase dynamics. Yet, they are typically limited by spatial resolution and remain challenging to implement with super-resolution microscopy approaches that require specific fluorophores or optical setups. Aurora kinase A (AURKA), a multifunctional serine/threonine kinase, has recently emerged as a critical regulator of mitochondrial physiology. However, visualising AURKA activation and activity in living cells with sub-diffraction precision remains a challenge.

Here, we introduce BioSenSRRF, a versatile approach that combines conventional FRET biosensors with Super-Resolution Radial Fluctuations (SRRF) microscopy. BioSenSRRF requires no modification of existing probes, can be implemented using standard microscopy setups, and is supported by a publicly available ImageJ framework to streamline image processing and data analysis. Using this pipeline, we imaged AURKA activation at mitochondria and its activity in regulating mitochondrial ATP levels. We uncover that AURKA activation and activity are compartmentalised into distinct mitochondrial domains containing the mitochondrial ATP synthase. These subdomains depend on the catalytic activity of AURKA, and they can be altered using previously validated AURKA inhibitors. Finally, we demonstrate that the cancer-associated polymorphism F31I enhances AURKA activation and ATP production on ATP synthase-enriched subdomains, underscoring its pathological relevance.

Altogether, BioSenSRRF provides a broadly accessible framework to enhance the spatial resolution of genetically encoded biosensors. This strategy opens new avenues for dissecting the subcellular organisation of protein complexes and their contribution to physiology and disease states.

## INTRODUCTION

Protein kinases are crucial players in the regulation of intracellular signalling across evolution ^1,2^. Before target phosphorylation, many kinases undergo an activation step through autophosphorylation. This step involves a three-dimensional conformational change of the catalytic cleft to accommodate the binding of a phosphate, and is required to prime the kinase for substrate recognition ^3–6^. Thanks to robust biochemical approaches, it is now established that a given kinase is virtually capable of phosphorylating a variety of substrates upon its activation ^6,7^. However, an intense area of investigation aims to uncover how each kinase targets only a subset of substrates at a given time, in a specific subcellular location, and in response to specific signals such as cell growth, division, metabolic regulation, or cell death.

Genetically encoded fluorescent biosensors have emerged as key tools to follow kinase activation and activity with spatiotemporal resolution ^8–12^. A majority of these sensors rely on Förster’s Resonance Energy Transfer (FRET) as a readout. A pair of fluorescent proteins with specific characteristics (i.e. spectral range, improved photostability, pH range, etc.) is used as a FRET donor/acceptor pair and constitutes the reporting unit of the biosensor ^10^. The full-length sequence of the kinase, or a substrate-specific peptide with a phospho-binding domain, are typical sensing units contained in biosensors. They indicate kinase activation through conformational changes or kinase activity, respectively ^10,12^.

Although there are over 100 genetically encoded kinase biosensors currently available, along with online tools compiling information on kinases, targets, fluorescent proteins, and readouts (https://biosensordb.ucsd.edu/), they only cover a fraction of the kinome ^2^. This gap is linked, among other factors, to the variety of kinase activation mechanisms ^5^ and to the large number and specificity of the substrates for each kinase ^9,13^. Current designs for substrate-based kinase biosensors require the *a priori* knowledge of the phosphorylation site(s), which are often substrate-specific. For proteins with 3D structures that are still under investigation, predicting phosphorylation sites can be challenging using *in silico* tools. Despite huge efforts to rationalise design strategies, biosensor backbones ^14–16^, the use of synthetic peptide libraries, and *in vitro* screening approaches ^17–20^, each new sensor requires extensive experimental validation to optimise its output and biological relevance. This creates a methodological bottleneck, hindering the development of probes for newly discovered substrates, understudied kinases, or kinase pools that are active in specific subcellular compartments. Overall, this leaves significant gaps in our capacity to track kinase activation and activity with adequate spatiotemporal resolution. This aspect is particularly relevant, as specific intracellular signaling actors were shown to be organised in micro- or nano-domains^21–23^.

Evidence that kinase-based signaling can also be organized at the nanoscale further highlights the need to increase the subcellular resolution of genetically encoded kinase biosensors ^24^. This can be done either by engineering new activation or kinase activity biosensors with improved localization properties or by increasing the spatiotemporal resolution of currently available probes. A variant of AKAR – a substrate biosensor specific for Protein Kinase A (PKA) – optimised for super-resolution imaging with photochromic stochastic optical fluctuation imaging (pcSOFI) highlighted the existence of plasma membrane microdomains where PKA is active ^24^. Contrary to the parental biosensor, which relies on FRET as a readout ^25^, the variant relies on the fluorescence fluctuation properties of the reporter unit. This behavior was called fluorescence Fluctuation INcrease by Contact (FLINC). In FLINC, the physical proximity between the two fluorescent proteins – Dronpa and TagRFP-T – induced by the activity of PKA increases the fluctuation of TagRFP-T, which can then be used to build high-resolution maps of PKA activity using pcSOFI. To date, FLINC can only be used with Dronpa and TagRFP-T, as no alternative fluorophore pair shows a similar fluctuation behavior.

Recent results show the cumulation of FRET with Super-resolution Optical Fluctuation Imaging (SOFI)^26^. SOFI is a super-resolution method that achieves a spatial resolution through the statistical analysis of multiple images of fluorescence from stochastically blinking fluorophores. The green photoswitchable fluorophore Skylan-S was used as the FRET donor, and the JF646 dye was used as the FRET acceptor. Through a theoretical model and the use of the FK506-binding protein (FKBP)/FKBP-rapamycin binding (FRB) domains of the mammalian target of rapamycin^27^, the authors super-resolve chemically-induced molecular interactions. Similarly to FLINC, however, this strategy requires specific donor/acceptor FRET pairs compatible with SOFI and the consequent modification of already-existing probes if they were to be used with this approach. In an independent study, the FKBP/FRB dimerization system has also been used to induce protein-protein interactions by adding rapamycin and by resolving them using Stimulated Emission Depletion (STED). The FKBP/FRB system was fused to the two moieties of the chemogenetic split reporter splitFAST2, and revealed with the HBR-3,5DOM fluorogenic chromophore^28^. Such a strategy requires the modification of constructs to insert the two halves of the splitFAST2 system, the addition of a compatible chromophore for STED-microscopy, and a STED imaging modality. Similarly, the use of specific fluorophore pairs and access to a STED device were key parameters in FRET-STED analyses performed in fixed samples^29^. Altogether, this body of evidence indicates that already-existing constructs should be re-engineered with pre-determined fluorophore pairs and revealed with specific imaging modalities to make them compatible with super-resolution techniques such as FLINC, SOFI, or STED.

In this study, we present a first innovative approach to increase the spatial resolution of genetically encoded biosensors by integrating FRET and super-resolution microscopy based on Super-Resolution Radial Fluctuations (SRRF) ^30,31^. We termed this approach BioSenSRRF. We show that BioSenSRRF can be used with currently available FRET biosensors and with standard acquisition and image analysis setups, without further adaptation. We also provide a publicly available, online analytical framework and a standalone software for BioSenSRRF image processing under the Fiji/ImageJ^32^ solution.

We here use this approach to correlate Aurora kinase A (AURKA) activation and activity at the mitochondria. Previous evidence has shown that AURKA is activated in this compartment, where it regulates ATP production by interacting with the mitochondrial respiratory chain ^33,34^. However, no substrate-specific biosensor is currently available to detect AURKA activity at mitochondria. By using a conformational biosensor detecting AURKA activation ^35^ and a second probe specific for mitochondrial ATP production (MitoGO-ATeam2) ^36^, we reveal AURKA activation and its direct regulation of ATP production at the nanoscale. With BioSenSRRF, we demonstrate that kinase activation and activity are organised into three classes of microdomains, which can be differentiated based on their FRET readout. We also discovered that currently available AURKA inhibitors only target selected classes of microdomains, but fail to completely inhibit kinase-dependent ATP production rates. Last, we identify that these microdomains correspond to clusters rich in the mitochondrial ATP synthase protein ATP5F1A. Cancer-related variants of AURKA induce an excessive ATP production at ATP5F1A-rich clusters, providing functional evidence for kinase activation and activity in mitochondrial microdomains.

## RESULTS

### Cumulative FRET and SRRF super-resolution microscopy improves the spatial resolution of a genetically encoded biosensor of kinase activation

Genetically encoded FRET biosensors are powerful tools to study the regulation of signaling pathways in cells ^8–12^. However, it can be problematic to determine the subcellular area where they are activated with sub-diffraction precision. Here, we combine previously characterized FRET biosensors ^35,37^ with super-resolution microscopy by SRRF to increase the spatial resolution of FRET events within mitochondria in cultured MCF7 cells. To obtain a complementary readout, we use a FRET biosensor reporting on kinase activation, and a second probe showing the consequences of kinase activity at mitochondria.

The AURKA biosensor performs FRET when the full-length kinase is activated through autophosphorylation on Threonine 288 ^35^. A conformational change brings the donor fluorophore (eGFP) fused on the N-terminus of the kinase, and the acceptor fluorophore (mCherry) fused on the C-terminus of AURKA at a FRET-permissive distance (< 10 nm) (**Figure 1A**). MCF7 cells expressing the AURKA biosensor and an acceptor-devoid control construct (eGFP-AURKA, hereby called Donor-only) under the control of the AURKA minimal promoter for endogenous-like expression were stained with MitoTracker DeepRed and imaged on a spinning-disk setup. For each channel, a total of 200 frames were acquired, and for each frame, the eGFP, mCherry, and MitoTracker channels were acquired sequentially. Then, images of the eGFP, mCherry and MitoTracker channels were reconstructed using SRRF post-acquisition methods (**Figure 1B**).

**Figure 1:**
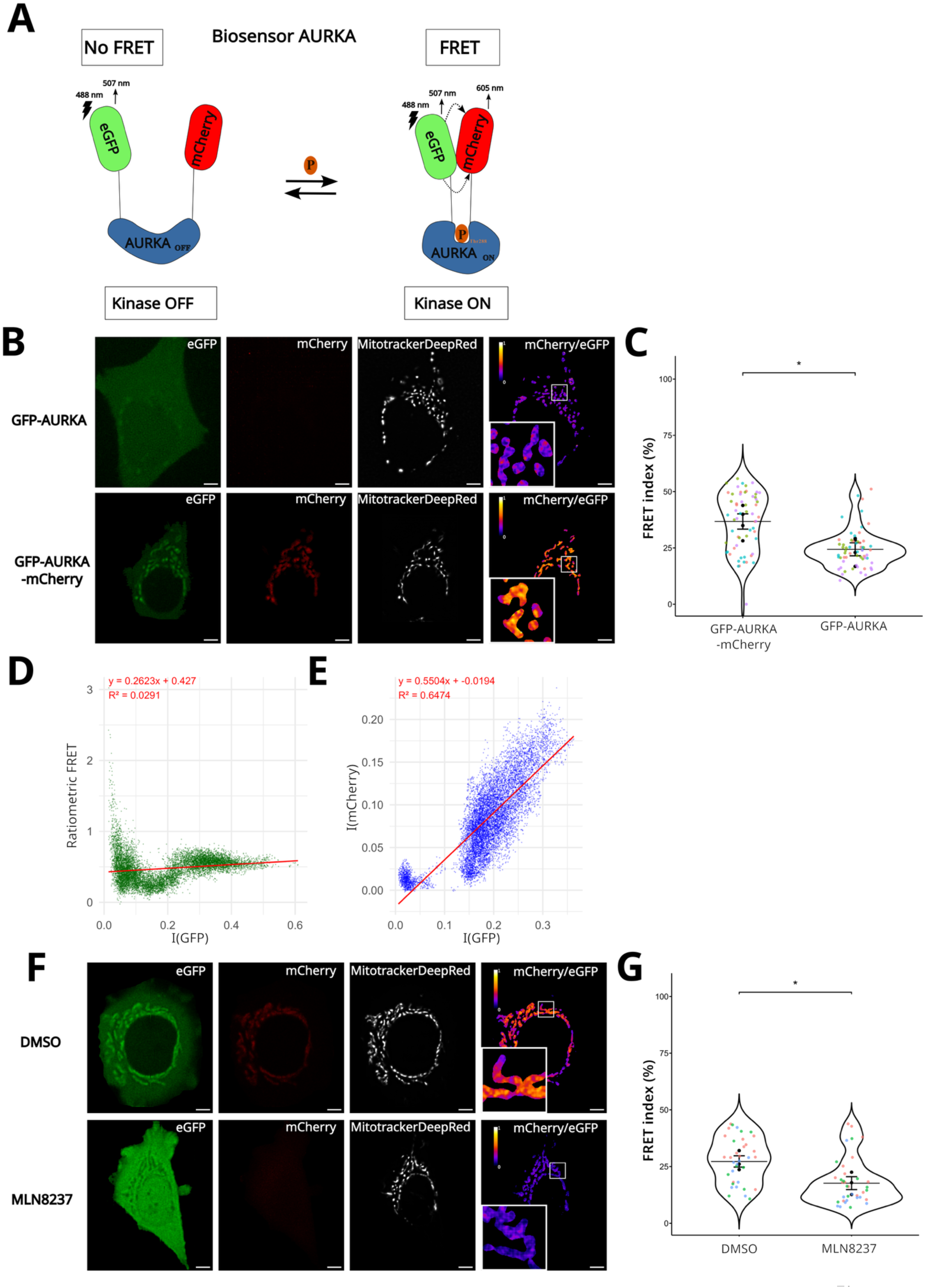
Validation of BioSenSRRF with the AURKA biosensor. **(A)** Cartoon showing the mode of functioning of the AURKA biosensor ^35^. The donor fluorophore is shown in green, the acceptor fluorophore is shown in red, and the entire kinase sequence is shown in blue. In the absence of kinase activation (Kinase OFF) the two fluorophores are apart and do not allow the transfer of energy by FRET (No FRET). When the kinase is autophosphorylated on T288 (Kinase ON), a conformational change of AURKA brings together its N- and C-termini, and it allows the two fluorophores to be at a FRET-permissive distance. **(B)** Super-resolved SRRF images of the GFP, mCherry (FRET), Mitotracker DeepRed channels, and visual representation of ratiometric FRET in cells expressing the control (Donor-only, upper panels) or the biosensor (lower panels). **(C)** SuperPlots of FRET Index values (0-100 %) on the mitochondrial area in cells expressing the biosensor or the donor-only counterpart. **(D-E)** Pixel-by-pixel correlation analyses on SRRF-reconstructed images between ratiometric FRET and GFP intensity (**D**), or between mCherry and GFP intensities (**E**). Linear regression equations and R² are indicated in red. **(F)** Super-resolved SRRF images of the GFP, mCherry (FRET), and Mitotracker DeepRed channels, and visual representation of ratiometric FRET in cells expressing the AURKA biosensor in cells treated with DMSO (upper panels) or the AURKA catalytic inhibitor MLN8237 (lower panels). **(G)** SuperPlots of FRET Index values (0-100 %) on the mitochondrial area in cells expressing the AURKA biosensor and treated with DMSO or MLN8237. Pseudocolour scale on FRET images: ratiometric FRET (0-1). Scale bar = 5 µm. Data are means ± SEM. Each dot in SuperPlots represents FRET index values from individual cells sorted by replicate (colours). *n* = 12 cells per condition in each independent replicate. Tukey test: \**P* < 0.05.

We first observed that, as previously reported, the mitochondrial network morphology was comparable between the two conditions. Then, the mitochondrial network between the two conditions was compared in terms of FRET index, which is an intensity ratio used relatively to a control condition (**Figure 1B-C**). The FRET index of the AURKA biosensor showed a mean of 36.73 % while that of the negative control was significantly lower (24.37 %). Due to the significant mean FRET index values of the donor-only construct, we wanted to exclude the possibility that SRRF could impact the FRET index *per se*. Therefore, we compared images before and after reconstruction. FRET index values before and after reconstruction showed a difference in the absolute values, with the biosensor showing a FRET index of 23.81 % and the donor-only showing a FRET index of 14.67 %. This could be due to the quality of the fluorescence signal corresponding to the MitoTracker channel in the raw images, which looks more diffused than in the reconstructed images. However, the difference in FRET index between the donor-only and the biosensor conditions was similar before and after reconstruction (**Figure S1A-B**). This highlights that SRRF enhances image resolution while maintaining net differences in FRET index between the biosensor and the control conditions. We then sought to verify whether the relative abundance of the biosensor in cells could affect the FRET index after reconstruction. The correlation between the FRET and the GFP signals was low (R²<0.5), indicating that the FRET index is not correlated with the amount of biosensor localised at mitochondria (**Figure 1D; Figure S2**). As expected, we also observed a strong correlation between the mCherry and eGFP intensity signals, which is due to the design of the biosensor (**Figure 1A, E**).

We then validated the readout of the BioSenSRRF pipeline to evaluate the effect of the catalytic AURKA inhibitor MLN2837 (Alisertib) ^38^ on the FRET index. After reconstruction, cells treated with MLN8237 showed a significantly lower FRET index (18.36 %) compared to those treated with the control compound DMSO (27.89 %) (**Figure 1F-G**). This is confirmed by a loss of the mCherry signal observed in cells treated with MLN8237 (**Figure 1F**), and it corroborates previous evidence on the capacity of this compound to abolish FRET within the AURKA biosensor.

To summarise, these results provide the first evidence that combining FRET and super-resolution by SRRF generates a pipeline capable of detecting the activation and the inhibition of the AURKA biosensor expressed at endogenous levels and located at mitochondria.

### The BioSenSRRF pipeline is compatible with a genetically-encoded biosensor for mitochondrial ATP production

As a further validation of the BioSenSRRF pipeline, we expressed the conformational biosensor MitoGO-ATeam2 to probe ATP production within mitochondria (**Figure 2A**). This biosensor is composed of a matrix-directed Mitochondrial Targeting Sequence (MTS), a circularly-permutated eGFP variant as the FRET donor, the ε subunit of the bacterial ATP synthase as the sensor part, and mKO2 as the FRET acceptor. The sensor undergoes a conformational change upon binding of ATP to the ε subunit of the ATP synthase. This brings the N and C termini of the probe in proximity for FRET. Similar to the AURKA biosensor, the donor-only MitoGFP-ATeam2 construct devoid of the acceptor was used as a negative control. Both MitoGFP-ATeam2 and MitoGO-ATeam2 show a mitochondrial localisation when transfected into MCF7 cells (**Figure 2B**). The calculation of the FRET index in SRRF-reconstructed images of the mitochondrial network showed that FRET is nearly absent in the donor-only condition (0.49 %). In comparison, the biosensor displays a mean FRET index of 43.74 % (**Figure 2C**).

**Figure 2:**
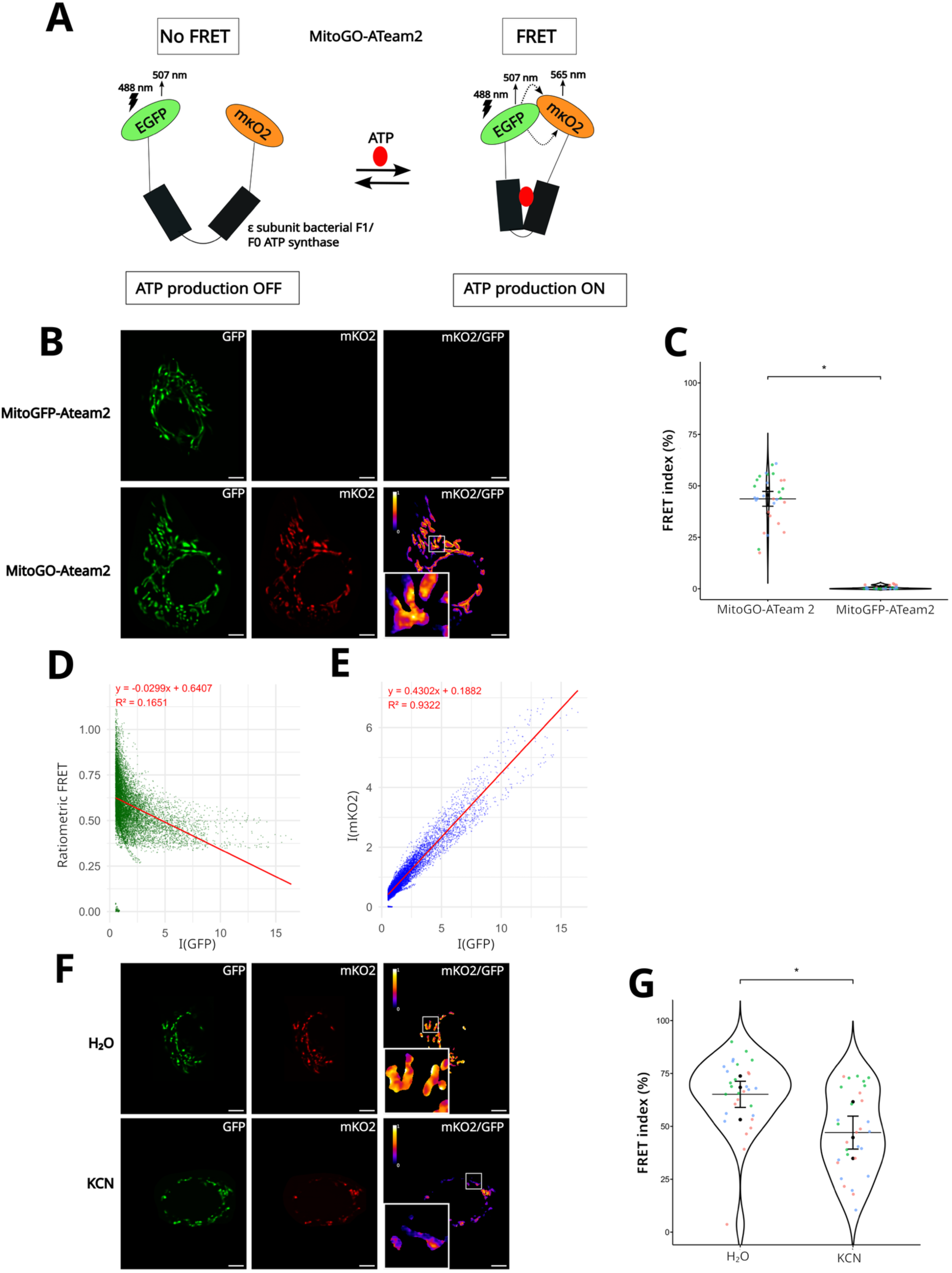
BioSenSRRF validation with the MitoGO-ATeam2. **(A)** Cartoon showing the mode of functioning of the MitoGO-ATeam2 biosensor ^36^. Donor fluorophore is shown in green (cpGFP), the acceptor fluorophore is shown in orange (mKO2), and the ε subunit of the bacterial ATP synthase is shown in black. The sensor is targeted to mitochondria thanks to a CoxVIII MTS (not shown). In the absence of ATP (ATP production OFF), the two fluorophores are apart and do not allow the transfer of energy by FRET (No FRET). In the presence of ATP (ATP production ON), a conformational change of the ε subunit brings together its N- and C-termini, allowing the two fluorophores to be at a FRET-permissive distance. **(B)** Super-resolved SRRF images of the GFP and mKO2 (FRET) channels, and visual representation of ratiometric FRET in cells expressing the control (Donor-only, upper panels) or the biosensor (lower panels). **(C)** SuperPlots of FRET Index values (0-100 %) in cells expressing the biosensor or the donor-only counterpart. **(D-E)** Pixel-by-pixel correlation analyses on SRRF-reconstructed images between ratiometric FRET and GFP intensity (**D**), or between mCherry and GFP intensities. (**E**) Linear regression equations and R² are indicated in red. **(F)** Super-resolved SRRF images of the GFP and mKO2 (FRET) channels, and visual representation of ratiometric FRET in cells expressing the MitoGO-ATeam2 biosensor treated with control (H2O, upper panels) or an ATP synthase inhibitor (KCN, lower panels). **(G)** SuperPlots of FRET Index values (0-100 %) in cells expressing MitoGO-ATeam2 and treated with H2O or KCN. Pseudocolour scale on FRET images: ratiometric FRET (0-1). Scale bar = 5 µm. Data are means ± SEM. Each dot in SuperPlots represents FRET index values from individual cells sorted by replicate (colours). *n* = 10 cells per condition in each independent replicate. Tukey test: \**P* < 0.05.

Similarly to what was observed with the AURKA biosensor, calculations performed on the raw images confirmed that SRRF reconstructions do not alter the FRET index (**Figure S1C**). Indeed, a FRET index of 51.23 % was obtained in cells expressing the MitoGO-ATeam2 biosensor while the signal in the donor-only condition showed a FRET index of 1.27 % (**Figure S1D**). For the MitoGO-ATeam2 biosensor as well, the correlation between mKO2 and eGFP intensities and that between FRET and eGFP signals indicated that FRET is not correlated with the expression level of the probe. We also verified that FRET is not influenced by photobleaching or diffusion of the probe within mitochondria on SRRF-reconstructed images (**Figure S2E-F; S3)**.

After verifying that the MitoGO-ATeam2 biosensor is compatible with the BioSenSRRF pipeline, we sought to reverse the FRET effect by treating cells with the mitochondrial Complex IV blocker potassium cyanide (KCN). Treatment with KCN is known to reduce the production of ATP within cells. Indeed, 1 hour of treatment with KCN was sufficient to show a significant reduction in the FRET index (47.06 %), compared to what was observed in control cells (65.14 %) (**Figure 2F-G**).

Our results confirm the previously-reported sensitivity of the biosensor to ATP inhibitors, and corroborate the usefulness of the BioSenSRRF pipeline to detect mitochondrial ATP availability under various conditions while increasing the localization precision of the biosensor.

### The random line analysis reveals spatially-resolved FRET Hotspots of ATP production

Previous studies have shown that AURKA requires an intact kinase activity for efficient mitochondrial localisation, as a kinase-dead form of the kinase (AURKA^K162M^) cannot be imported into mitochondria^33^. By using the BioSenSRRF pipeline, we confirmed that AURKA^K162M^ shows a lack of activity at mitochondria (**Figure S4**). The FRET index in cells overexpressing the kinase-dead version of the AURKA biosensor was significantly lower than that measured in cells expressing the non-mutated probe, confirming previous observations.

Previous biochemical analyses also revealed that AURKA overexpression increases mitochondrial ATP production^33^. However, it remains unexplored whether this increase is homogeneous or localised at specific sites within the mitochondrial network. To this end, we expressed the MitoGO-ATeam2 biosensor in the presence or absence of AURKA-iRFP670 or of AURKA^K162M^-iRFP670.

First, we observed an altered and condensed mitochondrial network in cells overexpressing AURKA compared to controls (**Figure 3A**). This is consistent with previous observations revealing that AURKA remodels mitochondrial morphology and induces the turnover of dysfunctional organelles^39^. Consistently, the comparison of FRET index means upon SRRF reconstruction revealed a statistically significant increase in cells overexpressing AURKA (61.39 %) compared to control cells (53.90 %), or cells expressing AURKA^K162M^ (49.38 %) (**Figure 3B**). This confirms that AURKA remodels mitochondrial ATP production in the organelles spared from degradation, and that AURKA kinase activity is required for this remodeling.

**Figure 3:**
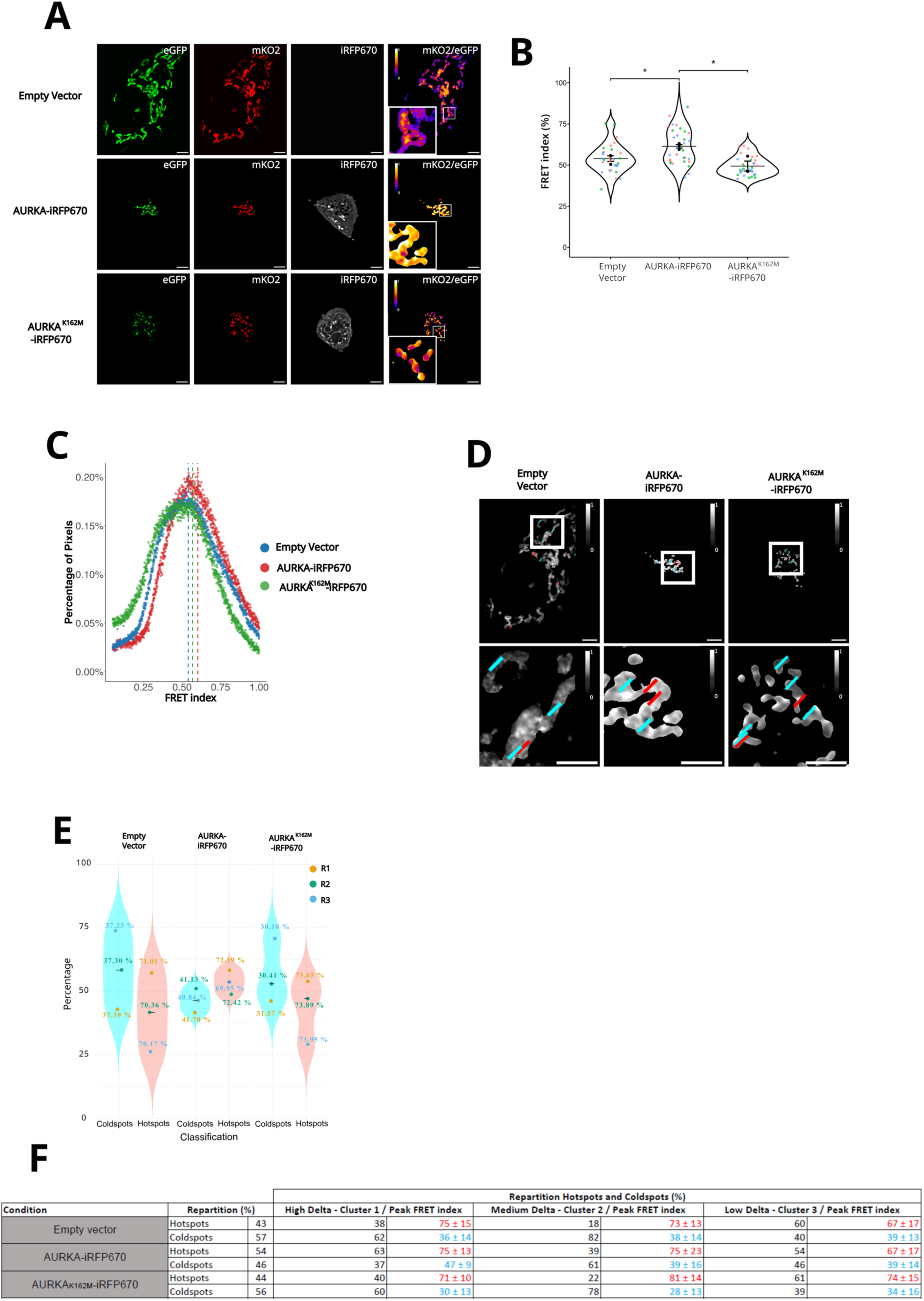
BioSenSRRF illustrates the spatial distribution of AURKA-dependent ATP production. **(A)** Super-resolved SRRF images of the GFP, mKO2 (FRET), and iRFP670 channels, and visual representation of ratiometric FRET in cells expressing MitoGO-ATeam2 together with an Empty vector (upper panels), AURKA-iRFP670 (middle panels), or AURKA^K162M^-iRFP670 (lower panels). **(B)** SuperPlots of FRET Index values (0-100 %) in the indicated conditions. **(C)** Histogram analysis showing the pixel-by-pixel distribution of FRET index values obtained from SRRF-resolved images of the MitoGO-ATeam2 biosensor in cells transfected as in (**A-B**). **(D)** Representative images of ratiometric FRET (shown in greyscale) upon random line analyses. Insets: higher magnification of the squared area. Scale bars: 5 µm (top images), 3 µm (insets). Red lines: line on a FRET hotspot; cyan lines: line on a FRET coldspot. **(E)** Violin plots representing the distribution of FRET Coldspots and Hotspots (in percentage) with the MitoGO-ATeam2 in the indicated conditions. Each dot represents the mean distribution values of each independent replicate (and Pseudocoloured in yellow, green, and blue), associated with its corresponding mean FRET index value. **(F)** Table showing the overall repartition of FRET Coldspots and Hotspots detected with the MitoGO-ATeam2 biosensor in the indicated transfection condition, together with the repartition of FRET Coldspots and Hotspots depending on the degree of variation of the FRET index (clusters). For each cluster and spot type, the FRET index peak value is shown in red (FRET Hotspots) or cyan (FRET Coldspots). *n* = 10 cells per condition in each independent replicate. Pseudocolour scale on FRET images: ratiometric FRET (0-1). Data are means ± SEM. Each dot in SuperPlots represents FRET index values from individual cells sorted by replicate (colours). One-way ANOVA followed by Tukey post-hoc test: \**P* < 0.05.

We then sought to determine the spatial localisation where ATP increase events take place. To this end, we used a pixel-by-pixel histogram analysis ^40^ to compare the distribution of FRET events within mitochondria in the three conditions. We first compared the histograms of FRET index values from cells in the three conditions (**Figure 3C**). We observed a shift in the histogram mode values of AURKA-overexpressing cells toward high FRET index values. The mode value of a histogram indicates the value with the highest frequency; a positive shift in the mode indicates that the distribution of FRET index values changes due to the presence of FRET events in the MitoGO-ATeam2 sensor. However, we also noticed that the shift is relatively modest, indicating that the overall distribution of FRET events is similar between the three conditions. This suggests that the increase in mean FRET index (**Figure 3B**) could be due to the existence of pools of highly-activated biosensor in discrete locations, rather than an overall increase in ATP production across the mitochondrial network.

To investigate the presence of clusters with high FRET index values within the mitochondrial network, we implemented a robust spatial analysis termed random line analysis. This analysis relies on 10 randomly drawn lines on the mitochondrial area, which were sufficient to cover the entire FRET variability of the mitochondrial network without line redundancy or overlap. For each line, we extracted the FRET index profiles, i.e. the pixel-by-pixel FRET index variations along the line (**Figure 3D**). Each line profile is categorised either as Gaussian (hereby FRET Hotspot) or inverse Gaussian (hereby FRET Coldspot) based on its discrete second derivative. We then used the silhouette coefficient method to assess the degree of similarity among line profiles. The score obtained with this approach determined that line profiles could be clustered into three distinct clusters using the k-means classification method. This method partitions data into k groups by minimising the distance between each point and the centroid of its associated cluster. Therefore, each line was assigned to a cluster depending on the variation of the FRET index along the line, hereby delta. These clusters emerged from the grouping generated by the k-means algorithm, allowing for an objective segmentation of the line profiles based on the extent of their FRET index variations. According to this classification, profiles showing a variation in the FRET index greater than 50 % between the minimum and maximum values were defined as having a high delta. Variations between 30 % and 50 % were classified as medium delta, while those below 30 % were considered low delta (**Figure S5**). We then explored the distribution of FRET Hotspots and Coldspots across all conditions, along with their corresponding mean FRET index values in three replicates (**Figure 3E**). Despite mean FRET index values being similar, our analysis revealed a higher percentage of FRET Hotspots in cells overexpressing AURKA compared to control cells, or cells expressing AURKA^K162M^ (**Figure 3E**). The random line analysis also revealed a lower percentage of FRET Coldspots in AURKA-overexpressing cells, compared to all other conditions (**Figure 3E**). Given these observations, we hypothesised that FRET Hotspots are more abundant in a cluster with high FRET variations – high delta – in AURKA-overexpressing cells. When looking at the distribution of FRET Hotspots and Coldspots in each independent cluster under these conditions, we observed that FRET Hotspots were more abundant in the cluster with high variations of the FRET index along the line (Cluster 1, High Delta), compared to all other conditions and to all other clusters (**Figure 3F**). This highlights that AURKA overexpression increases ATP production in specific spots within the mitochondrial network. The random line analysis also reveals no significant differences between cells transfected with an empty vector or cells overexpressing a kinase-dead version of AURKA, validating once again that kinase activity is required to increase mitochondrial ATP production.

Overall, the random line analysis deepens the readout of the BioSenSRRF pipeline by revealing Hotspots of ATP production within the mitochondrial network. Our results show that the kinase-dependent increase of mitochondrial ATP production is not evenly distributed, but it occurs at discrete locations that are spatially confined.

### Xanthohumol lowers AURKA activation and the abundance of AURKA-dependent ATP Hotspots

After describing FRET Hotspots as discrete sites of high metabolic activity and resulting from AURKA activation, we asked whether their abundance could be pharmacologically modulated.

Previous biochemical evidence has shown that Xanthohumol is a polyphenol with the capacity to rescue the metabolic rewiring induced by the overexpression of the kinase. However, it remains to be determined whether, by doing so, this compound also lowers AURKA activation. Therefore, we explored its mechanism of action at the subcellular level and with increased spatial resolution by evaluating its effect on AURKA activation, and on the mitochondrial activity of the kinase.

To determine the effect of Xanthohumol on AURKA activation, cells expressing the AURKA biosensor and stained with MitoTracker DeepRed were treated or not with this compound (**Figure 4A**). Cells treated with vehicle showed a mean FRET index of 51.25 %, while it was of 37.37 % in cells treated with Xanthohumol (**Figure 4B**). This significant difference in FRET index shows that Xanthohumol reduces the activation of AURKA within mitochondria. To study the effect of the drug on AURKA-dependent ATP production, we explored the readout of the MitoGO-ATeam2 biosensor in cells overexpressing AURKA or in controls treated with Xanthohumol. Mean FRET index calculations on SRRF-resolved images show that Xanthohumol does not affect the MitoGO-ATeam2 readout in cells that are not overexpressing AURKA. Indeed, control cells showed a mean FRET index of 52.49 % and 52.42 % in the presence and absence of Xanthohumol, respectively (**Figure 4C-D**). As expected, the mean FRET index in cells transfected with AURKA-iRFP670 and treated with vehicle was significantly higher (59.70 %) than in the previous conditions. On the contrary, treatment of AURKA-overexpressing cells with Xanthohumol significantly reduced the FRET index (48.33 %) (**Figure 4C-D**). These results confirmed previous biochemical analyses showing that Xanthohumol rescues ATP production in cells overexpressing AURKA.

**Figure 4:**
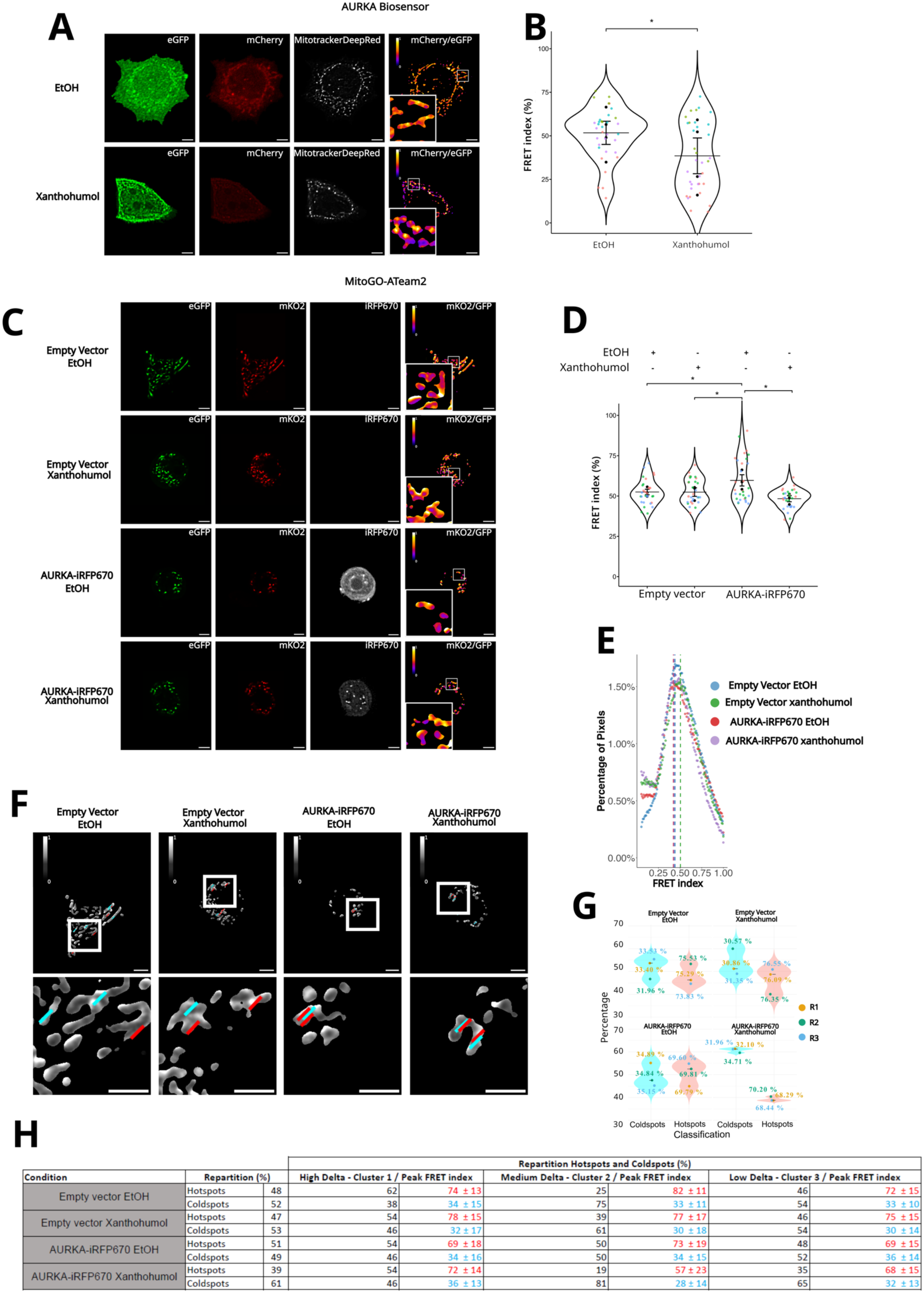
AURKA activation and activity are lowered in Xanthohumol-treated cells. **(A)** Super-resolved SRRF images of the GFP, mCherry (FRET), Mitotracker DeepRed channels, and visual representation of ratiometric FRET in cells expressing the AURKA biosensor in the presence of vehicle (EtOH, upper panels) or Xanthohumol (lower panels). **(B)** SuperPlots of FRET Index values (0-100 %) in cells expressing the AURKA biosensor and treated with EtOH or Xanthohumol. **(C)** Super-resolved SRRF images of the GFP, mKO2 (FRET) and iRFP670 channels, and visual representation of ratiometric FRET in cells co-expressing MitoGO-ATeam2 together with an Empty vector or AURKA-iRFP670, and treated with EtOH or Xanthohumol. **(D)** SuperPlots of FRET Index values (0-100 %) in cells expressing the MitoGO-ATeam2 biosensor and treated with EtOH or Xanthohumol. **(E)** Histogram analysis showing the pixel-by-pixel distribution of FRET index values obtained from SRRF-resolved images of the MitoGO-ATeam2 biosensor in cells transfected as in (**C-D**). **(F)** Representative images of ratiometric FRET (shown in greyscale) upon random line analyses. Insets: higher magnification of the squared area. Scale bars: 5 µm (top images), 3 µm (insets). Red lines: line on a FRET hotspot; cyan lines: line on a FRET coldspot. **(G)** Violin plots representing the distribution of FRET Coldspots and Hotspots (in percentage) with MitoGO-ATeam2 in the indicated transfection and treatment conditions. Each dot represents the mean distribution values of each independent replicate (and Pseudocoloured in yellow, green and blue), associated with its corresponding mean FRET index value. **(H)** Table showing the overall repartition of FRET Coldspots and Hotspots detected with the MitoGO-ATeam2 biosensor in the indicated transfection condition, together with the repartition of FRET Coldspots and Hotspots depending on the degree of variation of the FRET index (clusters). For each cluster and spot type, the FRET index peak value is shown in red (FRET Hotspots) or cyan (FRET Coldspots). *n* = 10 cells per condition in each independent replicate. Pseudocolour scale on FRET images: ratiometric FRET (0-1). Data are means ± SEM. Each dot in SuperPlots represents FRET index values from individual cells sorted by replicate (colours). Two-way ANOVA followed by Tukey post-hoc test: \**P* < 0.05.

After confirming previous evidence with imaging-based approaches, we asked whether Xanthohumol could reverse AURKA-dependent MitoGO-ATeam2 FRET Hotspots. After detecting no major differences in the pixel-by-pixel FRET distribution (**Figure 4E**), we performed random line analyses (**Figure 4F, Figure S6**), the silhouette clusterisation, and delta calculations (**Figure S6**) to identify FRET Hotspots and Coldspots. The random line analysis showed a dramatic reduction in the number of FRET Hotspots in cells overexpressing AURKA after treatment with the Xanthohumol (**Figure 4G)**. When looking at the repartition of FRET Hotspots and Coldspots into specific clusters, we observed that the Xanthohumol-dependent reduction of FRET Hostpots in AURKA-overexpressing cells impacts mostly the medium- and low-delta clusters **(Figure 4H, clusters 2 and 3)**. While FRET Hotspots in cluster 1 are not impacted by the drug and maintain a high level of ATP production, the amount of FRET Hotspots in clusters 2 and 3 is lowered. In addition, there is a concomitant increase in FRET Coldspots consistent with an overall decrease in the mean FRET index. Last, the Xanthohumol- dependent redistribution of FRET Hotspots and Coldspots remains unaltered in cells not overexpressing AURKA (**Figure 4G-H**).

Our data indicate that the number of AURKA-dependent FRET Hotspots can be pharmacologically modulated, and that Xanthohumol reduces their overall amount. Nevertheless, these results also show that Xanthohumol is specific to sites with medium or low variations of the FRET index, while leaving Hotspots with high FRET variations unaltered.

### ATP Hotspots are lowered with a mitochondrial-specific AURKA inhibitor

Given that Xanthohumol only alters the number of sites with medium or low MitoGO-ATeam2 FRET variations, we evaluated whether mitochondrial-specific inhibitors of AURKA could show increased specificity for FRET Hotspots with high FRET variations. Previous data showed that *N*-(4-hydroxy-3-methoxybenzyl)butyramide (HMBB), an analogue of the natural product Capsaicin, can restore AURKA-dependent turnover of defective mitochondria while leaving the kinase activated at centrosomes or in the nucleus ^41^. However, the effects of this compound on the activation of the mitochondrial pool of the kinase and on ATP production rates remain to be characterised.

We first explored the effect of HMBB on the activation of the AURKA biosensor at mitochondria. FRET index measurements in cells treated with HMBB show a lowered activation of the kinase (39.20 %) compared to cells treated with vehicle (47.11 %) (**Figure 5A-B**). This significant loss of FRET with HMBB indicates that the drug reduces the activation of the kinase specifically at mitochondria. This confirms previous structural predictions, which showed that the compound can bind to the kinase pocket of AURKA.

**Figure 5:**
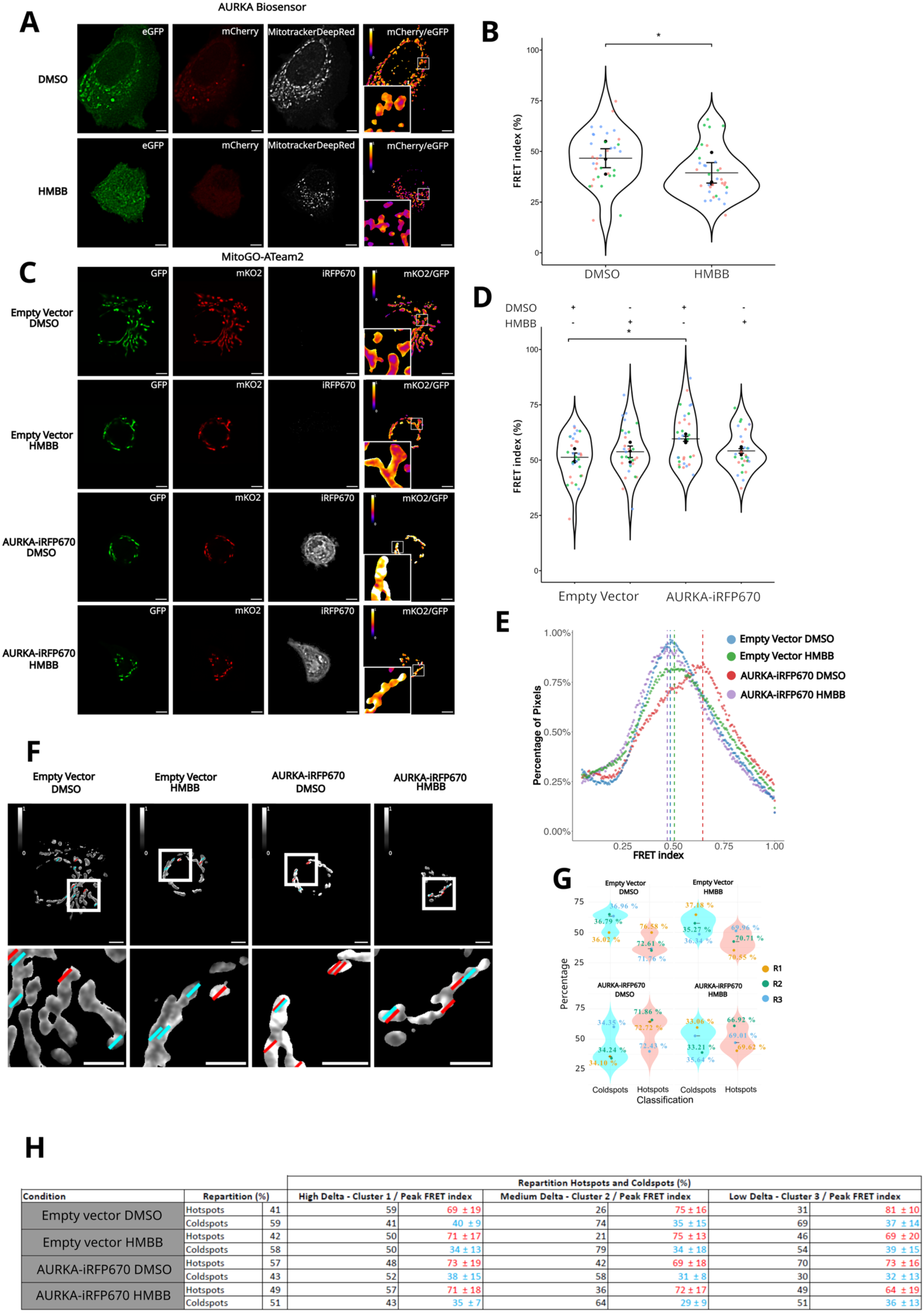
AURKA activation and activity are lowered in HMBB-treated cells. **(A)** Super-resolved SRRF images of the GFP, mCherry (FRET), Mitotracker DeepRed channels, and visual representation of ratiometric FRET in cells expressing the AURKA biosensor in the presence of vehicle (DMSO, upper panels) or HMBB (lower panels). **(B)** SuperPlots of FRET Index values (0-100 %) in cells expressing the AURKA biosensor and treated with DMSO or HMBB (*n* = 12). **(C)** Super-resolved SRRF images of the GFP, mKO2 (FRET), and iRFP670 channels, and visual representation of ratiometric FRET in cells co-expressing MitoGO-ATeam2 together with an Empty vector or AURKA-iRFP670, and treated with DMSO or HMBB. **(D)** SuperPlots of FRET Index values (0-100 %) in cells expressing the MitoGO-ATeam2 biosensor and treated with EtOH or Xanthohumol (*n* = 10). **(E)** Histogram analysis showing the pixel-by-pixel distribution of FRET index values obtained from SRRF-resolved images of the MitoGO-ATeam2 biosensor in cells transfected as in (**C-D**). **(F)** Representative images of ratiometric FRET (shown in greyscale) upon random line analyses. Insets: higher magnification of the squared area. Scale bars: 5 µm (top images), 3 µm (insets). Red lines: line on a FRET hotspot; cyan lines: line on a FRET coldspot. **(G)** Violin plots representing the distribution of FRET Coldspots and Hotspots (as a percentage) with MitoGO-ATeam2 in the indicated transfection and treatment conditions. Each dot represents the mean distribution values of each independent replicate (and Pseudocoloured in yellow, green, and blue), associated with its corresponding mean FRET index value. **(H)** Table showing the overall repartition of FRET Coldspots and Hotspots detected with the MitoGO-ATeam2 biosensor in the indicated transfection condition, together with the repartition of FRET Coldspots and Hotspots depending on the degree of variation of the FRET index (clusters). For each cluster and spot type, the FRET index peak value is shown in red (FRET Hotspots) or cyan (FRET Coldspots). *n* values refer to the number of cells per condition in each independent replicate. Pseudocolour scale on FRET images: ratiometric FRET (0-1). Data are means ± SEM. Each dot in SuperPlots represents FRET index values from individual cells sorted by replicate (colours). Tukey test: \**P* < 0.05 (**B**); Two-way ANOVA followed by Tukey post-hoc test: \**P* < 0.05 (**D**).

We then analysed the effect of HMBB on the FRET readout of MitoGO-ATeam2, in the presence or absence of overexpressed AURKA (**Figure 5C-D**). The mean FRET index of control cells with and without HMBB was 51.26 % and 53.75 %, respectively. This indicates that the drug has no effect on ATP production rates *per se*. Similar to previous analyses, the FRET index reached a mean of 59.61 % upon AURKA overexpression, while cells overexpressing AURKA and treated with HMBB showed a FRET index that decreased to 54.15 % (**Figure 5C-D**). Given that this mean decrease is overall modest, we analysed the SRRF-reconstructed images in a pixel-by-pixel manner.

Consistent with previous histogram analyses, we observed an increase in the FRET index distribution of cells overexpressing AURKA and treated with DMSO. This increase was abolished in cells overexpressing the kinase and treated with HMBB, indicating that the drug can reverse the kinase-dependent redistribution of FRET index values observed with the MitoGO-ATeam2 probe (**Figure 5E**). To investigate this redistribution at the spatial level, we performed a random line analysis (**Figure 5F; S6A-B**). The overall distribution of the mitochondrial FRET Hotspots observed in cells overexpressing AURKA was abolished upon treatment with HMBB (**Figure 5G**). When looking at the individual clusters, we observed that HMBB lowers the number of FRET Hotspots in clusters 2 and 3 (**Figure 5H**). Similar to what was observed in Xanthohumol-treated cells, FRET hotspots in cluster 1 are not impacted by the drug, and they show that MitoGO-ATeam2 biosensor is still active at these locations.

Overall, our results reveal a common mechanism of action for non-mitochondrial and mitochondrial-specific inhibitors of AURKA activity. The BioSenSRRF pipeline coupled to a random line analysis shows that both Xanthohumol and HMBB can lower the mitochondrial ATP production induced by an overexpression of the kinase. Despite these findings, our analyses also indicate that AURKA inhibition is cluster-specific and that both Xanthohumol and HMBB have no effect on mitochondria with high ATP production levels.

### The cancer-related AURKA polymorphism F31I alters AURKA activation and the spatial distribution of mitochondrial activity subdomains

Previous reports showed that the cancer-related AURKA polymorphism F31I confers susceptibility to several types of cancer ^42–44^. However, the functional consequences of this variant are poorly characterised, together with its role in AURKA-dependent ATP production. Since the overexpression of the F31I polymorphism results in more aggressive cancer phenotypes and low survival rates ^42–44^, we hypothesised that it could exacerbate the effect of AURKA on mitochondrial ATP levels.

Visually, MCF7 cells expressing AURKA or AURKA^F31I^ presented a reduced mitochondrial network, as expected (**Figure 6A**) ^45^. This observation was confirmed by mitochondrial mass measurements, where the mitochondrial volume of control cells was higher (15.49 %) than that of cells overexpressing AURKA or AURKA^F31I^ (10.70 % and 9.58 %, respectively) (**Figure 6B**). That implies that AURKA and AURKA^F31I^ reduce mitochondrial mass to a similar degree. Similarly, BioSenSRRF analyses of the MitoGO-ATeam2 FRET index normalised to the mitochondrial mass and histogram analyses revealed that AURKA and AURKA^F31I^ increase mitochondrial ATP levels to the same extent when compared to control cells (**Figure 6C**). Therefore, we asked whether the F31I polymorphism could alter the spatial distribution of mitochondrial FRET subdomains. To this end, we performed random line analyses on BioSenSRRF images (**Figure 6F; Figure S8**).

**Figure 6:**
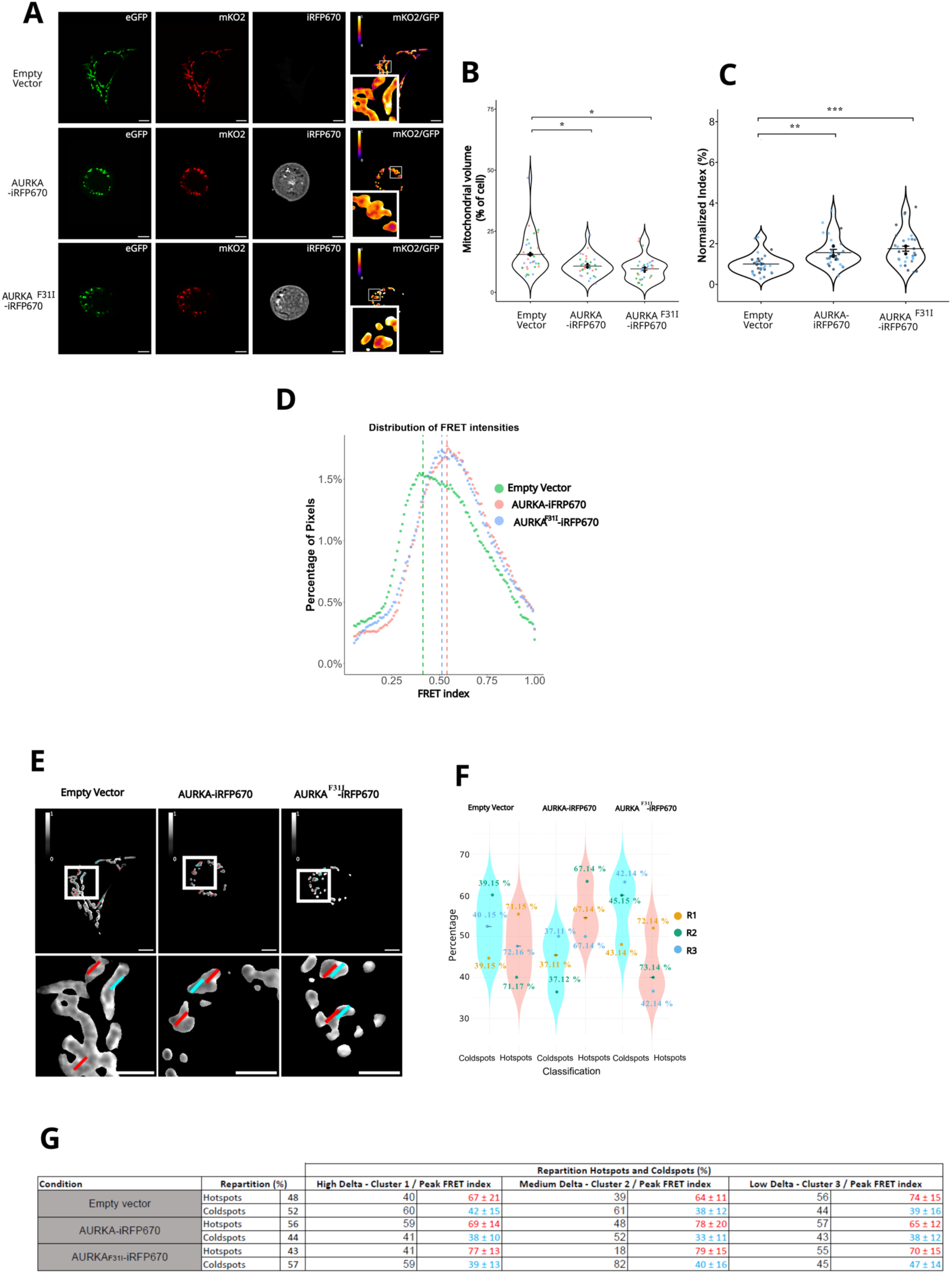
AURKA^F31I^ dramatically changes the distribution of FRET Hotspots and Coldspots within mitochondria. **(A)** Super-resolved SRRF images of the GFP, mKO2 (FRET) and iRFP670 channels, and visual representation of ratiometric FRET in cells expressing MitoGO-ATeam2 together with an Empty vector (upper panels), AURKA-iRFP670 (middle panels) or AURKA^F31I^-iRFP670 (lower panels). **(B)** SuperPlots of the mitochondrial area in the indicated conditions. **(C)** SuperPlots of FRET Index values (0-100 %) normalized against the mitochondrial area in the indicated transfection conditions. **(D)** Histogram analysis showing the pixel-by-pixel distribution of FRET index values obtained from SRRF-resolved images of the MitoGO-ATeam2 biosensor in cells transfected as in (**A-C**). **(E)** Representative images of ratiometric FRET (shown in greyscale) upon random line analyses. Insets: higher magnification of the squared area. Scale bars: 5 µm (top images), 3 µm (insets). Red lines: line on a FRET hotspot; cyan lines: line on a FRET coldspot. **(F)** Violin plots representing the distribution of FRET Coldspots and Hotspots (in percentage) with the MitoGO-ATeam2 in the indicated transfection conditions. Each dot represents the mean distribution values of each independent replicate (and Pseudocoloured in yellow, green, and blue), associated with its corresponding mean FRET index value. **(G)** Table showing the overall repartition of FRET Coldspots and Hotspots detected with the MitoGO-ATeam2 biosensor in the indicated transfection conditions, together with the repartition of FRET Coldspots and Hotspots depending on the degree of variation of the FRET index (clusters). For each cluster and spot type, the FRET index peak value is shown in red (FRET Hotspots) or cyan (FRET Coldspots). Pseudocolour scale on FRET images: ratiometric FRET (0-1). Data are means ± SEM. Each dot in SuperPlots represents FRET index values from individual cells sorted by replicate (colours). *n* = 10 cells per condition in each independent replicate. One-way ANOVA followed by Tukey post-hoc test: \**P* < 0.05, \*\**P* < 0.01, \*\*\**P* < 0.001.

As shown above, the distribution of FRET hotspots and coldspots in cells overexpressing AURKA showed a majority of FRET Hotspots (56 %) when compared to control cells (48%). Surprisingly, cells overexpressing AURKA^F31I^ showed an opposite FRET behavior. Under these conditions, the distribution of FRET hotspots lowered more than what was observed in control cells (43 %) (**Figure 6G-H**). When looking at the number of Hotspots and Coldspots in each cluster, we observed that the F31I polymorphism dramatically lowered the number of FRET Hotspots in clusters 1 and 2 while showing a concomitant rise in FRET Coldspots compared to cells overexpressing AURKA (**Figure 6H**). Overall, these data show that there is a global increase in ATP within mitochondria of cells expressing AURKA^F31I^, but also more submitochondrial domains where ATP levels are low (FRET Coldspots).

While these data indicate that the F31I polymorphism changes the spatial distribution of ATP within the organelle, they also raise a conundrum. How can global ATP levels rise, while FRET hotspots decrease? To answer this question, we verified whether this was a result of an altered AURKA activation at mitochondria in the presence of a F31I variant. We compared the mitochondrial FRET readout of the AURKA biosensor in the presence or in the absence of the F31I variant using the BioSenSRRF pipeline. Our results showed no significant differences in the mean mitochondrial activation of the AURKA^F31I^ biosensor compared to the AURKA biosensor without the polymorphism (mean FRET index of 33.11 % and 32.94 %, respectively) (**Figure 7A-B**). These observations were confirmed by similar results with the histogram analysis (**Figure 7C**). Random line analyses and spot distribution analyses also showed similarities between AURKA and AURKA^F31I^ **Figure 7D-F; Figure S9**). However, we noticed a greater variability in the overall distribution of FRET hotspots in cells overexpressing the F31I variant (**Figure 7E**), which was supported by a reduction of the frequencies of cluster 3 (low delta) compared to those in cells expressing normal AURKA (72 % to 81 %, respectively).

**Figure 7:**
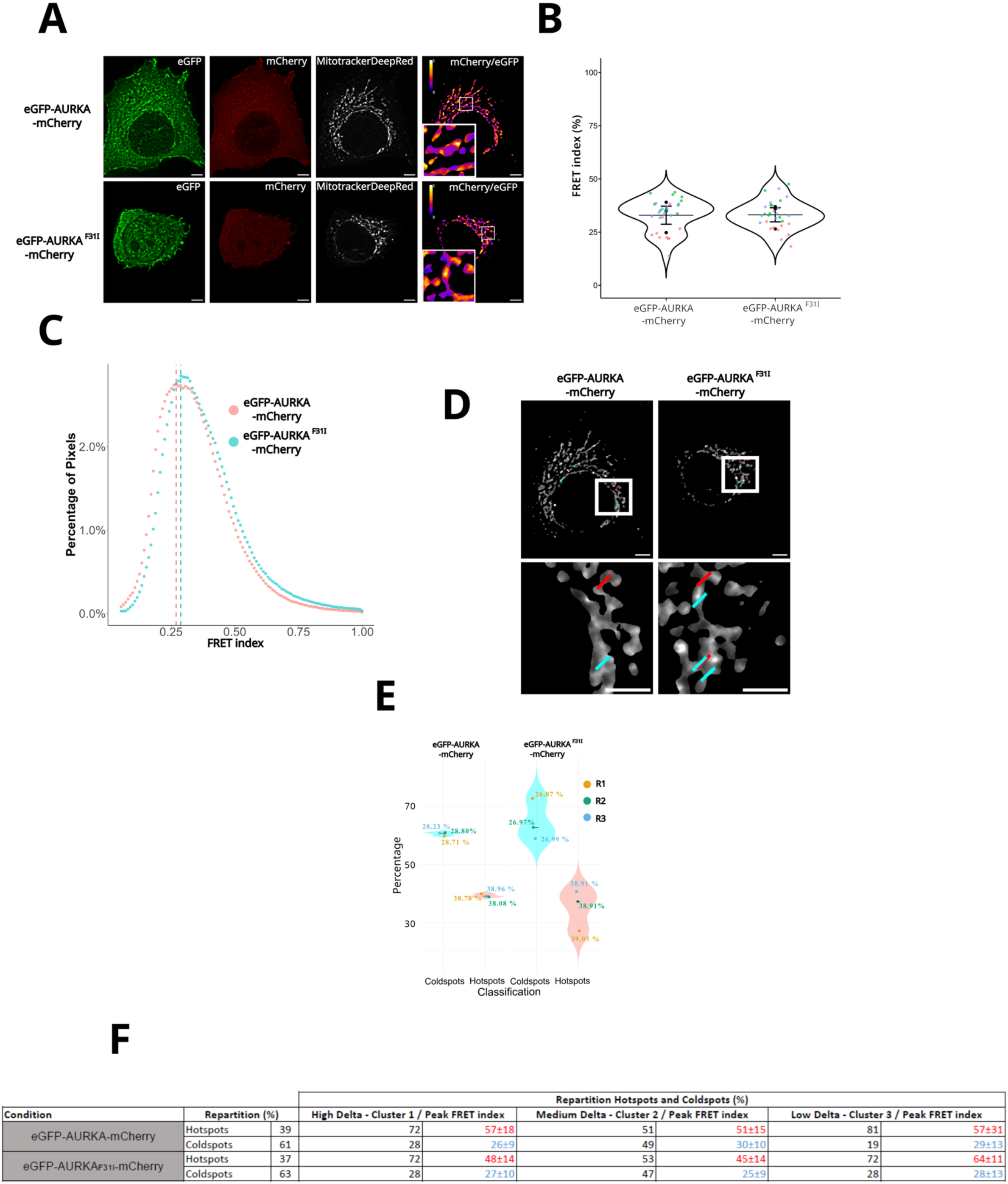
AURKA^F31I^ mildly alters AURKA activation within mitochondria. **(A)** Super-resolved SRRF images of the GFP, mCherry (FRET), and Mitotracker DeepRed channels, and visual representation of ratiometric FRET in cells expressing the AURKA (upper panels), or the AURKA^F31I^ biosensors (lower panels). **(B)** SuperPlots of FRET Index values (0-100 %) in the indicated conditions. **(C)** Histogram analysis showing the pixel-by-pixel distribution of FRET index values obtained from SRRF-resolved images of the indicated AURKA biosensors, as in (**A-B**). **(D)** Representative images of ratiometric FRET (shown in greyscale) on the MitoTracker DeepRed-positive area upon random line analyses. Insets: higher magnification of the squared area. Scale bars: 5 µm (top images), 3 µm (insets). Red lines: line on a FRET hotspot; cyan lines: line on a FRET Coldspot. **(E)** Violin plots representing the distribution of FRET Coldspots and Hotspots (in percentage) with the indicated AURKA biosensors in the indicated transfection conditions. Each dot represents the mean distribution values of each independent replicate (and Pseudocoloured in yellow, green, and blue), associated with its corresponding mean FRET index value. **(F)** Table showing the overall repartition of FRET Coldspots and Hotspots detected with the indicated AURKA biosensors, together with the repartition of FRET Coldspots and Hotspots depending on the degree of variation of the FRET index (clusters). For each cluster and spot type, the FRET index peak value is showed in red (FRET Hotspots) or cyan (FRET Coldspots). Pseudocolour scale on FRET images: ratiometric FRET (0-1). Data are means ± SEM. Each dot in SuperPlots represents FRET index values from individual cells sorted by replicate (colours). *n* = 10 cells per condition in each independent replicate. Tukey test: comparisons were not significant.

Overall, BioSenSRRF reveals significant differences in the spatial distribution of AURKA^F31I^ submitochondrial activity domains, which are correlated with differences in its activation.

### BioSenSRRF reveals AURKA activation and activity on the ATP synthase

Since F31 is contained in the AURKA MTS ^33^, we asked whether the increase in activation and activity of AURKA^F31I^ could be explained by its differential abundance inside mitochondria. We thus verified the effect of the F31I polymorphism on the import of the protein into mitochondria by performing Western blot analyses and the quantification of the total (46 kDa) and mitochondrial isoforms of the protein (43 and 38 kDa)^33^ (**Figure 8A-B**). We observed an increased abundance of the mitochondrial AURKA43 and AURKA38 isoforms in cells expressing the AURKA^F31I^ mutant compared to those in cells expressing normal AURKA. This shows a higher mitochondrial import of AURKA^F31I^ compared to the normal protein. To support this evidence, we performed molecular 3D models of the MTS of AURKA, the F31I variant, and of the MTS of Aurora kinase A of 2 other mammals that constitutively possess an isoleucine at a conserved position (**Figure 8C-D; Figure S10**). These modelling results indicated that the F31I mutation in the MTS of the human AURKA increases the flexibility of the C-terminal part of the α-helix. Interestingly, this is not observed in the MTS of Aurora kinase A in mouse and panda, where the isoleucine does not increase the flexibility of the elbow after the α-helix. The increase in flexibility observed with the F31I variant could support a facilitated import of the kinase through the mitochondrial membranes, resulting in higher AURKA amounts undergoing activation and activity.

**Figure 8:**
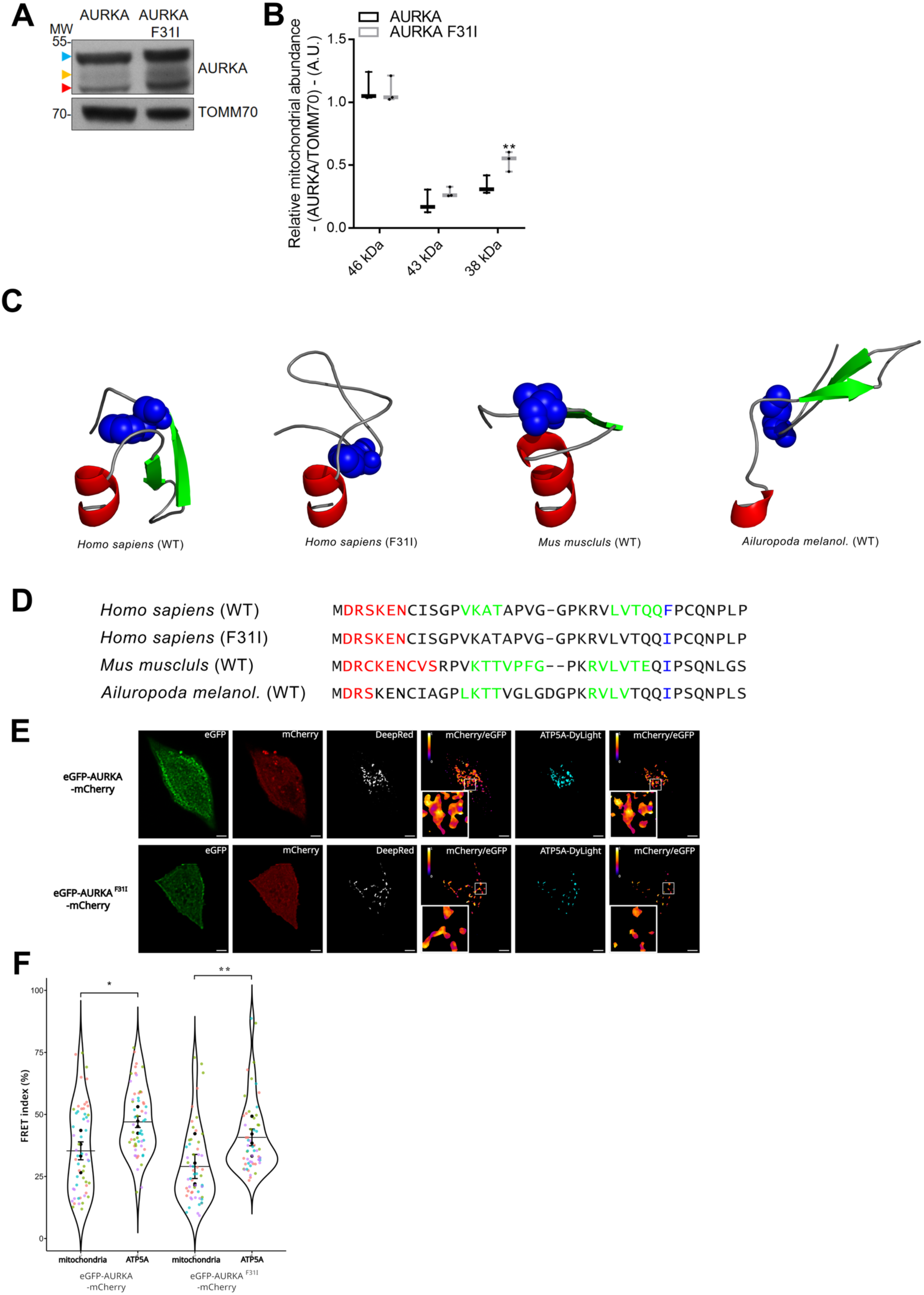
BioSenSRRF reveals the activation of AURKA on mitochondrial Complex V. **(A)** Western blotting analyses of mitochondrial fractions blotted against AURKA and TOMM70 (used as loading control) in HEK293 cells expressing AURKA-6xHis or AURKAF31I-6xHis. The colored arrows indicate the total AURKA isoform, and the two mitochondrially-processed isoforms ^33^ (blue: 46 kDa, yellow: 43 KDa, Red: 36 kDa). **(B)** Boxplot of the mitochondrial abundance of the three isoforms of AURKA relative to that of TOMM70. *n* = 3 independent experiments. **(C)** Top 1 structural models of the MTS peptide of AURKA located at its N-terminal end (amino acids 1-38) computed by PEP-Fold4. Normal AURKA, AURKA^F31I^, and the MTS of mouse and panda AURKA homologs are shown. Green: β-sheet; red: α-helix; blue spheres: Phenylalanine or Isoleucine residues; grey: coils. **(D)** Sequence alignment of the four MTSs illustrated in (**C**). Green: β-sheet; Red: α-helix; blue spheres: Phenylalanine or Isoleucine; black: coils. **(E)** Super-resolved SRRF images of the GFP, mCherry (FRET), Mitotracker DeepRed, and the ATP5A-DyLight 405 channels. Visual representation of ratiometric FRET is shown for the MitoTracker and the ATP5A channels (Ratiometric FRET panels from left to right) in cells expressing the normal or the F31I biosensors. **(F)** SuperPlots of FRET Index values (0-100 %) on the MitoTracker DeepRed- or on the ATP5A-positive areas in cells expressing the normal or the F31I biosensors. Pseudocolour scale on FRET images: ratiometric FRET (0-1). Data are means ± SEM. Each dot in SuperPlots represents FRET index values from individual cells sorted by replicate (colours). *n* = 14 cells per condition in each independent replicate. Two-way ANOVA and the Holm-Sidak method (**B**), Two-way ANOVA followed by Tukey post-hoc test (**F**): \**P* < 0.05, \*\**P* < 0.01, \*\*\**P* < 0.001.

Since AURKA^F31I^ is more abundant than AURKA within mitochondria, we hypothesized that its activation and activity would be further concentrated at specific intramitochondrial sites. This concentration would thus change the spatial distribution of FRET Hotspots and Coldspots across the entire mitochondrial network, as previously observed (**Figure 6**). To this end, we expressed the AURKA or AURKA^F31I^ biosensor in MCF7 cells and monitored its activation directly at sites containing the ATP synthase. We used ATP5A as a proxy of the ATP synthase, and as an interactor of AURKA within this complex previously identified with FRET/FLIM analyses ^34^. This protein was labelled with an anti-ATP5A primary antibody and decorated with a Dylight 405 secondary antibody. We then used the BioSenSRRF pipeline to extract FRET index values at sites positive for ATP5A, and compared them with FRET index values calculated on the entire mitochondrial area (**Figure 8E)**. As expected, the mean FRET index in mitochondria did not change between AURKA and AURKA^F31I^ (35 % and 30 %, respectively), indicating similar activation. However, the mean FRET index on the ATP5A-positive area significantly increased in cells expressing AURKA, indicating that the kinase preferentially activates at ATP5A-positive sites (**Figure 8F**). Furthermore, this increase is stronger in cells expressing the F31I variant. This indicates that the mutated kinase is significantly more active on ATP synthase-positive clusters than in the overall mitochondrial area.

In conclusion, we provide evidence that BioSenSRRF increases the spatial resolution of genetically encoded FRET biosensors of kinase activation and activity. With this pipeline, we uncover clusters of AURKA activation and of ATP production within mitochondria. We show that the kinase activity of AURKA regulates these clusters and that AURKA-specific inhibitors have only partial effects on them. Using BioSenSRRF, we find that these clusters are enriched in Complex V and that AURKA is activated at these submitochondrial domains. These results showcase the generalisability of BioSenSRRF towards a multitude of genetically encoded probes, and its potential to track biochemical activities at the nanoscale.

## DISCUSSION

The sensitivity of BioSenSRRF reveals the existence of mitochondrial subdomains with metabolic relevance. The power of FRET reveals kinase activation and activity using genetically encoded biosensors ^35,36^, and combining them with SRRF-based super-resolution microscopy ^30,46^ further enhances the spatial resolution of these events with sub-diffraction precision. BioSenSRRF has two main advantages. First, its implementation can be done on standard microscopy setups, without the need to purchase or adapt already-existing equipment. In principle, as long as fluctuations can be observed, a variety of setups and illuminations can be used. The *parameter sweep* function of the native eSRRF plugin ^46^ allows users to verify whether acquisitions meet the standards for image reconstruction in super-resolution mode. After determining the best reconstruction parameters, our automated BioSenSRRF pipeline will first calculate FRET on SRRF images (https://github.com/cyclochondria/BioSenSRRF.git) (**Figure 9**). A second part of the pipeline is dedicated to data analysis, enabling end users to transition from raw images to statistics without requiring coding (**Figure 9**). The code can be customised by more experienced users, allowing for the implementation of complementary data analysis methods. Alternatively, a ready-to-use version of the pipeline is available as a standalone software for a quick application of BioSenSRRF (<u>10.5281/zenodo.17312923</u>)^47^. Second, BioSenSRRF relies on already available biosensors, with fluorophores compatible with the vast majority of existing microscopy setups. Indeed, kinase sensors often undergo cycles of optimisation to increase their dynamic range, adapt fluorescent proteins to specific needs (e.g., increased brightness, resistance to pH variations, avoiding dimerisation, etc.), or to use them with specific microscopy approaches ^9,10^. The constant upgrades of kinase sensor features are an area of intense investigation. However, novel sensor variants often require a re-validation of the probe to ensure that its output and sensitivity to activators or inhibitors are preserved. Since this re-validation is often cumbersome, we believe that BioSenSRRF will fuel discoveries in cell biology thanks to its applicability to previously engineered FRET probes undergoing conformational changes.

**Figure 9:**
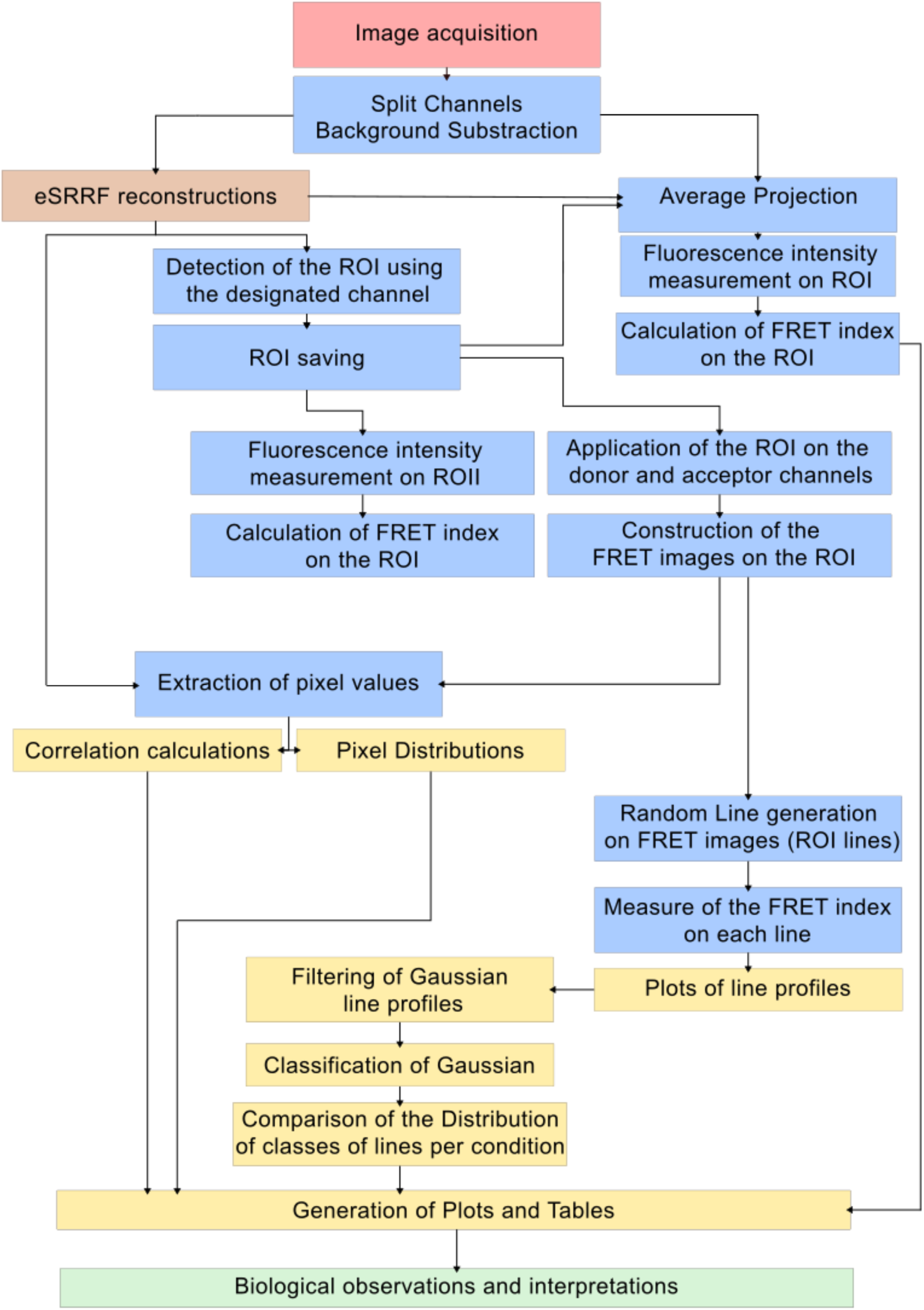
Workflow of the BioSenSRRF pipeline. Visual representation of the key steps of the pipeline. The colors of boxes are as follows: pink, image acquisition; blue, image processing and extraction of pixel intensities; brown, eSRRF Super-Resolution processing; yellow: calculations, statistical analyses, and plot generation; green: interpretations of the results by the user.

With 200 frames collected for each fluorescent channel (∼200 msec per frame) and with the acquisition setup described here, tracking analyses revealed either absent or minimal diffusion of the probes within mitochondria, along with no significant photobleaching events. While temporal and spatial resolutions will certainly be improved by faster cameras in the near future, the current temporal resolution might fall short for tracking extremely rapid events. This is due to the global acquisition time required to image the three channels. However, the total number of frames is a parameter that can be optimised by the end user, and a lower number of frames could be a pertinent choice if a higher temporal resolution is required. Another key parameter for optimising the readout of BioSenSRRF is the estimation of the background signal, so that it can be reduced before the SRRF reconstruction. Since FRET is calculated ratiometrically, a high background in the donor and/or the acceptor channels would give rise to artefacts ^48^. Background determination is a parameter that needs to be experimentally determined and subtracted by the end user. Background values should be optimised and standardized for each probe, their expression levels, the setup, and the imaging parameters used during acquisition ^48^.

Although we demonstrated that SRRF is a convenient super-resolution approach to further increase the spatial resolution of the FRET readout, it should be kept in mind that the FRET Hotspots and Coldspots inevitably include the reporting and sensing units of each biosensor. While the probes would still accurately reflect changes in their environment, such as kinase activation and changes in ATP levels in our case, the current BioSenSRRF configuration may still induce a slight increase in the absolute size of FRET Hotspots and Coldspots detected by the random line analyses (**Figure S5**). A potential solution would be that donor and acceptor moieties are inserted *at locus*, and that the exogenous expression of the entire sensor is avoided. However, it should also be considered that a vast majority of FRET-based biosensors are chimeric proteins or protein fragments. Their genomic insertion would be unfeasible or hard to achieve without having complex cellular phenotypes arising. Even if the *at locus* insertion were possible, mitochondria present a further layer of complexity. Proteins located on the Outer and Inner Mitochondrial Membranes are often localised in clusters, and this includes the complexes of the mitochondrial respiratory chain ^49,50^. Imaging modalities such as dSTORM are suitable to determine the dimensions of each cluster, although they are often not compatible with living samples. Altogether, BioSenSRRF allows for a spatial resolution higher than conventional imaging and is compatible with live cell imaging. FRET index values can be calculated on super-resolved images and compared relatively across experiments. We also demonstrate that its readout can be further enhanced when using protein-specific antibodies, which further refine the spatial readout of FRET under fixed conditions.

With BioSenSRRF, we correlated kinase activation and activity in the absence of a substrate biosensor. Our strategy turned out to be particularly suitable when the substrate of the kinase or the site of phosphorylation is unknown. By using an activity sensor with a broader readout, such as ATP levels, BioSenSRRF reports on the kinase activity of AURKA that may be exerted through multiple phosphorylations, either on the same or on different substrates located in proximity. While substrate-based biosensors would be instrumental in dissecting the precise post-translational modifications at play, our data show that function-based readouts are also convenient for monitoring organelle-specific AURKA activity. We identified three clusters according to the extent of FRET variations with the MitoGO-ATeam2 sensor. In light of recent discoveries showing metabolic heterogeneity within a single mitochondrial network ^51^, it is tempting to speculate that the three clusters may be a readout of this functional specialisation. Although our data show that AURKA and AURKA^F31I^ both activate on ATP5A-positive clusters, this activation pattern seems to be predominant for AURKA^F31I^ (**Figure 8**). The partial overlap between AURKA activation Hotspots and ATP5A raises the interesting possibility that other proteins may be contained in these Hotspots, and act as preferential partners of AURKA for activation. This could be similar to the AURKA-TPX2 interaction on the mitotic spindle ^52–55^. In this light, BioSenSRRF is a convenient strategy to uncover structural domains at the nanoscale and to identify the proteins located at these domains. This approach could easily be extended to probes reporting on mitochondrial signal transduction pathways ^56^, to proteins regulating mitochondrial ultrastructural stability, and to mtDNA. BioSenSRRF could also be used to screen for novel inhibitors that bind to and inhibit AURKA activity when in proximity to respiratory chain complexes. Since AURKA is a kinase with compartmentalised activations and activities – including at mitochondria ^33,34,39,45^, at centrosomes ^57–61^, and at the nucleus ^62^, other kinases could likely adopt specific intraorganellar activation and activity patterns. The spatiotemporal and mechanistic regulation of these patterns can be revealed by BioSenSRRF-based imaging of existing or novel fluorescent probes.

## MATERIALS AND METHODS

### Expression vectors and molecular cloning

DNA constructs were generated using Gibson Assembly Master Mix (New England Biolabs) or with T4 DNA ligase (Thermo Fisher Scientific). All restriction enzymes were purchased from Thermo Fisher Scientific. All cloning reactions were verified on a 3130 XL sequencer (Applied Biosystems). All site-directed mutagenesis reactions were performed using QuikChange site-directed mutagenesis (Agilent Technologies). The complete list of plasmids used in the study is reported in Supplementary Table 1.

### Cell culture, expression vectors and chemicals

MCF7 (HTB-22) and HEK293 (CRL-1573) human cell lines were purchased from the American Type Culture Collection (ATCC) and grown in Dulbecco’s modified Eagle’s medium (DMEM, Sigma-Aldrich) supplemented with 10 % fetal bovine serum (FBS, GE Healthcare), 1% L-glutamine (GE Healthcare), and 1 % penicillin-streptomycin (GE Healthcare) at 37°C. eGFP-AURKA and eGFP-AURKA-mCherry cells were generated by transfecting MCF7 cells with the corresponding mammalian expression vector by the *AURKA* minimal promoter sequence (CTTCCGG) as in ^35^ and in the presence of the X-tremeGENE HP transfection reagent (Roche), following the manufacturer’s instructions. Stable clones were selected in DMEM supplemented with 10 % FBS, 1 % L-glutamine, 1 % penicillin–streptomycin and 500 mg ml ^-1^ geneticin (Sigma-Aldrich). Cells were grown in Nunc Lab-Tek Chambers (Thermo Fisher Scientific), or in polymer-coated µ-Slide 8 Well high slides (Ibidi) for live imaging. Cells were grown on glass coverslips placed in 24-well plates (Falcon) for immunofluorescence experiments, or on 10-cm^2^ Petri dishes for Western blotting. Plasmid transfections were performed with Lipofectamine 2000 (Thermo Fisher Scientific) according to the manufacturer’s instructions. Cells were imaged or fixed 48 hours post transfection. For a list of plasmids used in this study, see **Table 1**. Before proceeding with live cell imaging, standard growth media was replaced with phenol red-free Leibovitz’s L-15 medium (Thermo Fisher Scientific) supplemented with 20 % FBS and 1 % penicillin–streptomycin.

Cells were treated with MLN8237 (SelleckChem) at a final concentration of 100 nM for 3 hours before imaging. Xanthohumol (SelleckChem) was used at a final concentration of 30 µM and incubated for 24 hours before fixation, while HMBB was used at a final concentration of 10 µM for 24 hours before fixation or live cell imaging. Potassium cyanide (KCN, Sigma-Aldrich) was used at a final concentration of 300 µM for 1 hour before live cell imaging.

### Mitochondrial dyes, cell fixation and immunostaining

MitoTracker DeepRed (Thermo Fisher Scientific) was used to stain mitochondria at a final concentration of 20 nM. Cells were incubated with the dye for 30 minutes at 37°C in complete growth media and washed twice with complete growth media before fixation or live cell imaging.

Cells were fixed in 4% paraformaldehyde (Sigma-Aldrich), permeabilised with 0.2% Triton X-100, saturated with 5% BSA, and stained using standard immunocytochemical procedures. The A monoclonal anti-ATP5A (clone 7H10BD4F9) antibody (Invitrogen) was used at a 1:500 dilution. An anti-mouse DyLight 405 secondary antibody (Thermo Fisher Scientific) was used at a 1:500 dilution. Cells were then mounted in ProLong Gold antifade reagent (Thermo Fisher Scientific) before imaging.

### Ratiometric FRET imaging

Images were acquired using an inverted microscope equipped with a spinning disk head (Leica) coupled to an EMCCD camera, with a 63x oil immersion objective, and controlled by the Inscoper software (Inscoper, France). The excitation and emission wavelengths for donors (eGFP, cpGFP) were 488 and 525/40 nm; a 488 nm excitation and a 607/36 nm emission for acceptors (mCherry/mKO2); a 638 nm excitation and 685/40 nm for MitoTracker DeepRed and iRFP670, and a 405 nm excitation and 447/60 nm for DyLight 405. For each channel, 200 frames were acquired with an exposure time of 25 ms to image the donor and the acceptor of MitoGO-ATeam2, and 50 ms to image the donor and the acceptor of the AURKA biosensor. The exposure time for Mitotracker DeepRed, iRFP670-containing constructs and Dylight 405 was 50 ms.

### Image treatment, FRET index calculation and SRRF-based reconstructions

The raw images were processed using ImageJ/Fiji ^32^ with a publicly available custom-made macro (https://github.com/cyclochondria/BioSenSRRF.git). The macro is divided in multiple steps to facilitate image treatment. First, each raw image undergoes background subtraction. To this end, images are thresholded to delete pixels with intensity below 600 gray levels for AURKA, or 800 gray levels for the MitoGO-ATeam2 biosensors, respectively. Images are then reconstructed with the eSRRF plugin ^46^ using settings predetermined with the *parameter sweep* function. In our case, the following settings were used: *Magnification=5, Radius=3, Sensitivity=3, Number of frames=200, Average reconstruction*. These parameters gave the best *Quality and Resolve factor (*based on *FRC* and *RSP)* on the images. The mitochondrial area was determined using the auto-threshold function in Li mode on the MitoTracker DeepRed channel for the AURKA biosensor, or the GFP channel for the MitoGO-ATeam2 biosensor. The mitochondrial area positive for ATP5A was determined with the auto-threshold function in MaxEntropy mode on the DyLight 405 channel. The FRET index before or after SRRF reconstruction was calculated by dividing the fluorescence intensity on the acceptor channel (mCherry/mKO2) by the intensity of the donor channel (GFP/cpGFP) on the MitoTracker DeepRed-positive area for the AURKA biosensor, the GFP-positive area for the MitoGO-ATeam2 biosensor, or on the DyLight 405-positive area for ATP5A. FRET images were produced using the *Image Calculator* tool embedded within Fiji. Z-projections were generated using the Fiji *Z-projection* tool on the 200 raw images acquired for each channel. The Fiji plug-in *Trackmate* ^63,64^ was used to follow the signal on the entire stack of raw images, to ensure the absence of probe diffusion over time.

### Correlation and histogram analyses

Intensity correlation analyses were performed after extracting pixels from SRRF images. The correlations were calculated using the RStudio software with a publicly available custom-made script (https://github.com/cyclochondria/BioSenSRRF.git). 10,000 random pixels were used for each correlation analysis. R² values were then used to determine the degree of correlation between channels. Histogram analyses were also performed after extracting pixels from SRRF images, and processed with a second, custom-made RStudio script to calculate the FRET index distribution (https://github.com/cyclochondria/BioSenSRRF.git). This feature is also present in the standalone BioSenSRRF software ^47^.

### Random line analysis

The random line analysis was performed on FRET images after SRRF reconstruction using Fiji (https://github.com/cyclochondria/BioSenSRRF.git). Each line was set at a dimension of 25 pixels and is inclined at 45°. Due to the mitochondrial area being smaller in cells expressing the AURKA^F31I^ variant, lines were set at a dimension of 15 pixels for experiments in **Figures 7** and **8**. For each image, 10 random lines were generated, and the FRET index on each line was then calculated. The lines were filtered and classified in clusters based on the variation of FRET index values from min to max, and analyzed along the line. The optimal number of clusters was determined using the Silhouette coefficient, an indicator used to assess the internal consistency of the formed clusters. This index ranges from -1 to 1, with a value above 0.5 indicating a good separation between groups. To strengthen the reliability of this approach, the number of iterations required for convergence within each cluster was also analysed. The distribution of delta FRET index values was performed using the k-means algorithm, an unsupervised classification method partitioning data into k distinct groups. The value of k was 3 throughout the datasets (**Figure S5-S9**). This approach then defines stable groups by reassigning points to the nearest centroids and adjusting the centroids until they stabilise. The optimal number of clusters was calculated through the Silhouette coefficient based on the ΔFRET values. This approach identifies natural groups within the data without imposing manual thresholds, thereby avoiding potential bias that could arise from an arbitrary segmentation, and is independent of the raw intensity FRET values of the biosensors. The lines showing a Gaussian and an Inverse Gaussian profile were then extracted according to their curvature calculated through their discrete second derivative, and the percentage of distribution of Gaussian (FRET Hotspots) and Inverse Gaussian (FRET Coldspots) were plotted for each replicate. The distribution of FRET Hotspots and Coldspots was then classified according to the variation of FRET along the line (High, medium and low variations) for each replicate. The random line generation and filtration pipeline was generated on RStudio (https://github.com/cyclochondria/BioSenSRRF.git) and adapted for the standalone software^47^.

### Molecular modelling

As no homology sequences were found for the N-terminal sequences of AURKA in the Protein Data Base (PDB), molecular models were produced using the PEP-fold method ^65^ through a dedicated web server (version 4). According to the confidence level of the software, the top 5 models were retained. The models were then colored according to the secondary structure observed, and the Isoleucine/Phenylalanine residues were shown as blue spheres.

### Mitochondrial fractionation and Western blotting

Isolated mitochondrial fractions were obtained by differential centrifugation as previously described ^66^. Protein concentrations were assayed using Bradford reagent (Bio-Rad) and boiled in Laemmli buffer, separated with SDS-PAGE and transferred onto nitrocellulose membrane (GE Healthcare). The primary antibodies used were a primary monoclonal mouse anti-AURKA (Clone 5C3 ^67^) and anti-TOMM70 (Abcam, Ab106193) used at a 1:20 and 1:5000 dilution, respectively. An anti-mouse, horseradish-peroxidase conjugated secondary antibody was purchased from Jackson ImmunoResearch Laboratories. Membranes were incubated with a commercially available chemiluminescence reagent (Pierce). Chemiluminescence was captured on CP-BU new films and a CURIX 60 developer (AGFA HealthCare). The relative abundance of AURKA import bands was calculated by normalizing their intensity toward that of the loading control (TOMM70).

### Statistical analyses

To compare FRET index on Super-plots, Tukey tests were performed with RStudio. The comparison of AURKA isoforms abundance was calculated using two-way ANOVA and the Holm-Sidak method. Linear regression equations were determined on RStudio using R². The calculation of K-means and the silhouette score were calculated using RStudio, and ideal clusterization was determined using the K-means graphics. The statistical tests performed are reported in each Figure legend.

## Supporting information

Jolivet, Desmaison et al Supp Figures

## ACKNOWLEDGEMENTS

The authors wish to thank Ricardo Henriques and Hannah S. Heil from the Henriques team (ITQB Nova) for helpful advice on the SRRF implementation and for critical reading of the manuscript. We thank Malvina Salami, Anaïs Le Guerer, and Maiwenn Cherré for technical assistance. HMBB was kindly provided by Laurent Désaubry (Univ. Strasbourg, France). We thank Stéphanie Dutertre at the Microscopy Rennes Imaging Center (MRic, BIOSIT, Biogenouest) for assistance. MRic is a member of the national infrastructure France-BioImaging, supported by the French National Research Agency (*ANR-24-INBS-0005 FBI BIOGEN*). We thank all the members of our team for constructive discussions.

## AUTHOR CONTRIBUTIONS

CRediT taxonomy: Conceptualization (G.B.), Data Curation (N.J, P-J.D, X.P., O.D., G.B.), Formal Analysis (N.J, P-J.D, X.P.,O.D., G.B.), Funding Acquisition (N.J., G.B.), Investigation (N.J, P-J.D, X.P., O.D., G.B.), Methodology (N.J, P-J.D, X.P., O.D., G.B.), Project Administration (G.B.), Resources (G.B.), Software (N.J, P-J.D, A.M., X.P., O.D., G.B.), Supervision (G.B.), Validation (G.B.), Visualization (N.J, P-J.D, O.D., G.B.), Writing – original, review and editing (N.J, P-J.D, X.P., O.D., G.B.).

## DATA AVAILABILITY

Key source microscopy data and standalone versions of the BioSenSRRF software are available on Zenodo^47^ (DOI: <u>10.5281/zenodo.17312923</u>). Fiji/ImageJ and RStudio custom-made macros for BioSenSRRF analyses are available on https://github.com/cyclochondria/BioSenSRRF.git. All other data are available from the corresponding author (G.B.) upon request.

