## Supplementary material for "Combining FRET and super-resolution microscopy reveals kinase activation and mitochondrial activity at the nanoscale": Jolivet, Desmaison et al Supp Figures

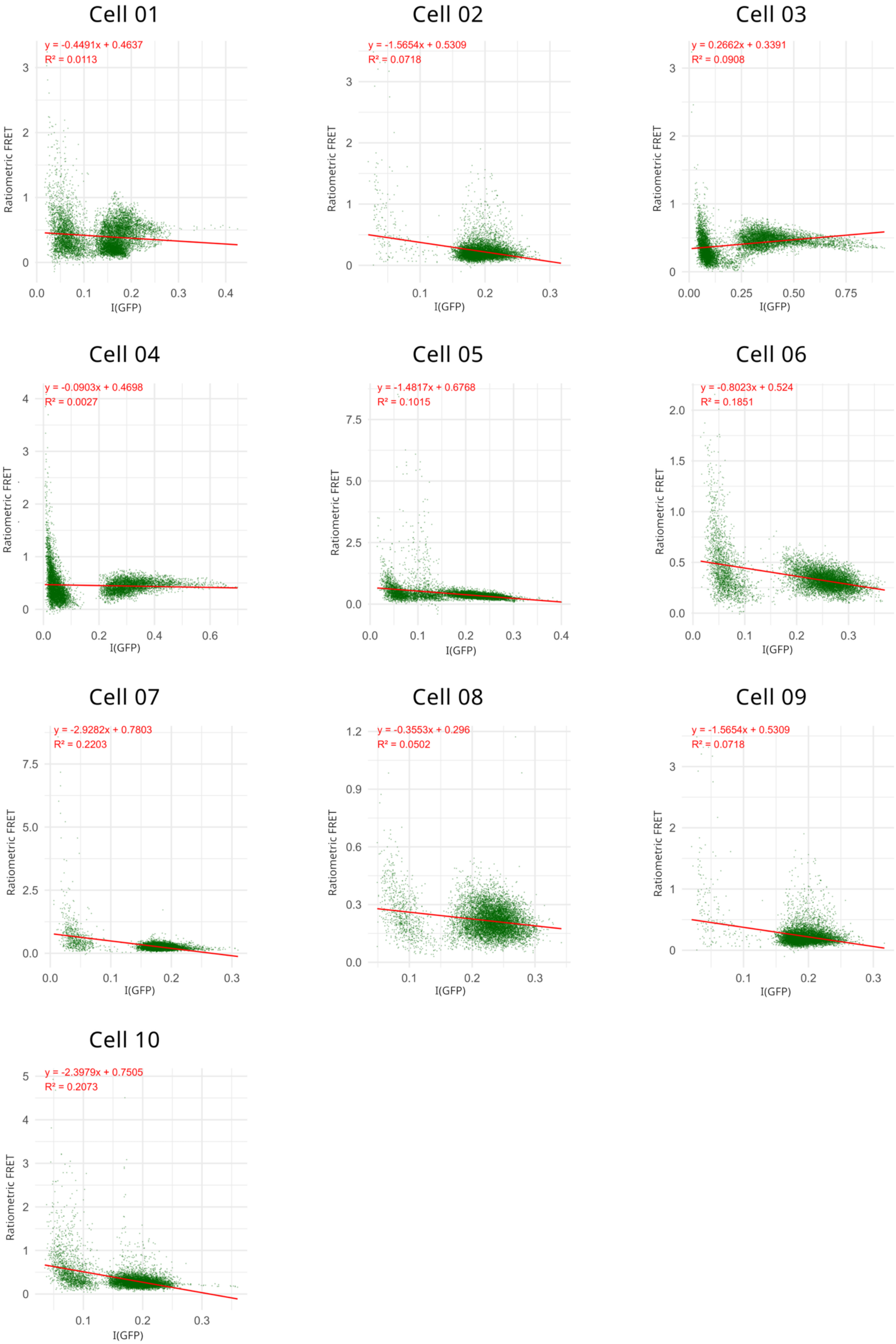
Cell 01
y = -0.4491x + 0.4637
$R^2$ = 0.0113
Ratiometric FRET
I(GFP)
Cell 02
y = -1.5654x + 0.5309
$R^2$ = 0.0718
Ratiometric FRET
I(GFP)
Cell 03
y = 0.2662x + 0.3391
$R^2$ = 0.0908
Ratiometric FRET
I(GFP)
Cell 04
y = -0.0903x + 0.4698
$R^2$ = 0.0027
Ratiometric FRET
I(GFP)
Cell 05
y = -1.4817x + 0.6768
$R^2$ = 0.1015
Ratiometric FRET
I(GFP)
Cell 06
y = -0.8023x + 0.524
$R^2$ = 0.1851
Ratiometric FRET
I(GFP)
Cell 07
y = -2.9282x + 0.7803
$R^2$ = 0.2203
Ratiometric FRET
I(GFP)
Cell 08
y = -0.3553x + 0.296
$R^2$ = 0.0502
Ratiometric FRET
I(GFP)
Cell 09
y = -1.5654x + 0.5309
$R^2$ = 0.0718
Ratiometric FRET
I(GFP)
Cell 10
y = -2.3979x + 0.7505
$R^2$ = 0.2073
Ratiometric FRET
I(GFP)

**Figure S1: Linear regression analyses on independent cells expressing the AURKA biosensor**

Linear regression analyses showing the absence of regression between FRET index and GFP intensities (A.U.) for 10 individual cells expressing the AURKA Biosensor. Linear regression equations and $R^2$ are indicated in red.

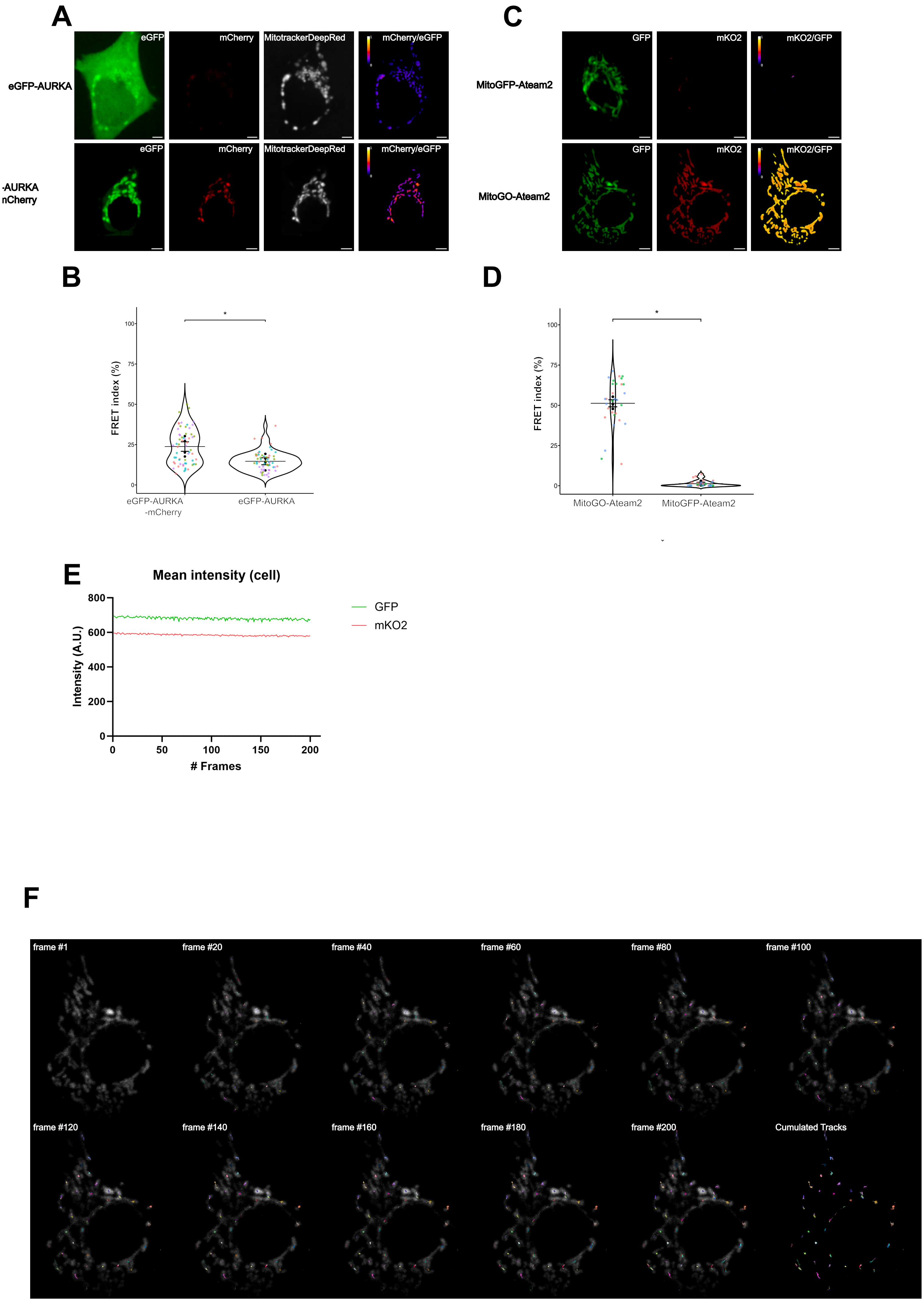

A
eGFP
mCherry
MitotrackerDeepRed
mCherry/eGFP
eGFP-AURKA
-AURKA
nCherry
C
GFP
mKO2
mKO2/GFP
MitoGFP-Ateam2
MitoGO-Ateam2
B
FRET index (%)
eGFP-AURKA
-mCherry
eGFP-AURKA
D
MitoGO-Ateam2
MitoGFP-Ateam2
E
Mean intensity (cell)
Intensity (A.U.)
### Frames
GFP
mKO2
F
frame #1
frame #20
frame #40
frame #60
frame #80
frame #100
frame #120
frame #140
frame #160
frame #180
frame #200
Cumulated Tracks

**Figure S2: FRET measurement on raw-z projection**

**(A)** Average projections of raw image stacks (200 frames each) of the GFP, mCherry (FRET) and Mitotracker DeepRed channels, and visual representation of ratiometric FRET in cells expressing the control (Donor-only, upper panels) or the AURKA biosensor (lower panels). **(B)** SuperPlots of FRET Index values (0-100 %) on the mitochondrial area in cells from (**A**) expressing the biosensor or the donor-only counterpart. **(C)** Average projections of raw image stacks (200 frames each) of the GFP and mKO2 (FRET) channels, and visual representation of ratiometric FRET in cells expressing the control (Donor-only, upper panels) or the biosensor (lower panels). **(D)** SuperPlots of FRET Index values (0-100 %) in cells from (**C**) expressing the MitoGO-ATeam2 biosensor or the donor-only counterpart. **(E)** Representative mean intensity (A.U.) measured over the 200 frames in the GFP and mKO2 channels on cells as in (C). **(F)** Representative timepoints of GFP signal tracking over the 200 frames. The colored lines indicate the center of each tracked object over the frames. The last image shows all the tracks detected along the 200-frame acquisition. $n$ = 10 cells per condition in each independent replicate. Pseudocolour scale on FRET images: ratiometric FRET (0-1). Data are means ± SEM. Each dot in SuperPlots represents FRET index values from individual cells sorted by replicate (colours). Tukey test: $^*P < 0.05$.

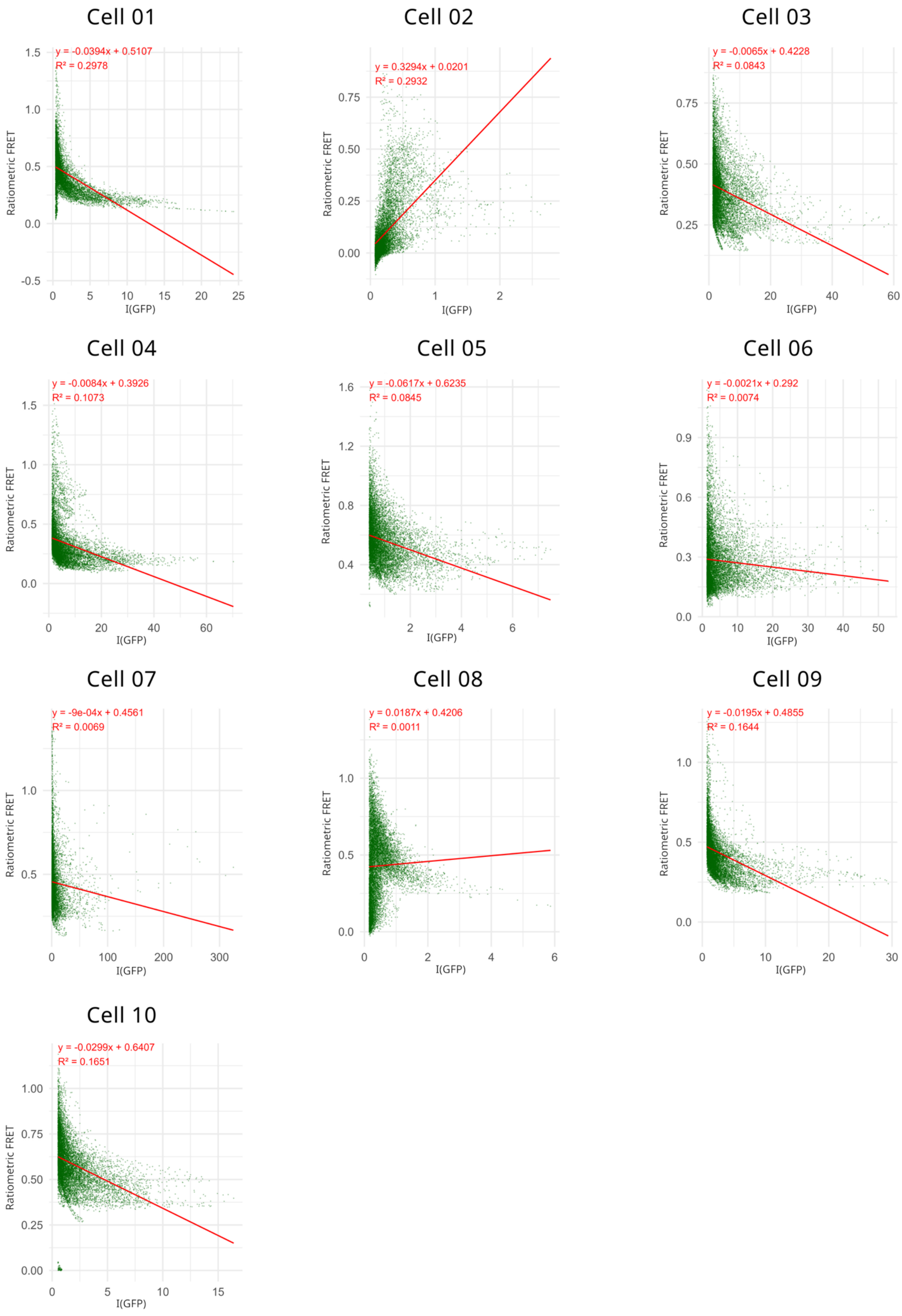

Cell 01
y = -0.0394x + 0.5107
R² = 0.2978
Ratiometric FRET
I(GFP)
Cell 02
y = 0.3294x + 0.0201
R² = 0.2932
Ratiometric FRET
I(GFP)
Cell 03
y = -0.0065x + 0.4228
R² = 0.0843
Ratiometric FRET
I(GFP)
Cell 04
y = -0.0084x + 0.3926
R² = 0.1073
Ratiometric FRET
I(GFP)
Cell 05
y = -0.0617x + 0.6235
R² = 0.0845
Ratiometric FRET
I(GFP)
Cell 06
y = -0.0021x + 0.292
R² = 0.0074
Ratiometric FRET
I(GFP)
Cell 07
y = -9e-04x + 0.4561
R² = 0.0069
Ratiometric FRET
I(GFP)
Cell 08
y = 0.0187x + 0.4206
R² = 0.0011
Ratiometric FRET
I(GFP)
Cell 09
y = -0.0195x + 0.4855
R² = 0.1644
Ratiometric FRET
I(GFP)
Cell 10
y = -0.0299x + 0.6407
R² = 0.1651
Ratiometric FRET
I(GFP)

**Figure S3: Linear regression analyses on independent cells expressing the MitoGO-ATeam2 biosensor**

Linear regression analyses showing the absence of regression between FRET index and GFP intensities (A.U.) for 10 individual cells expressing the MitoGO-ATeam2 Biosensor. Linear regression equations and $R^2$ are indicated in red.

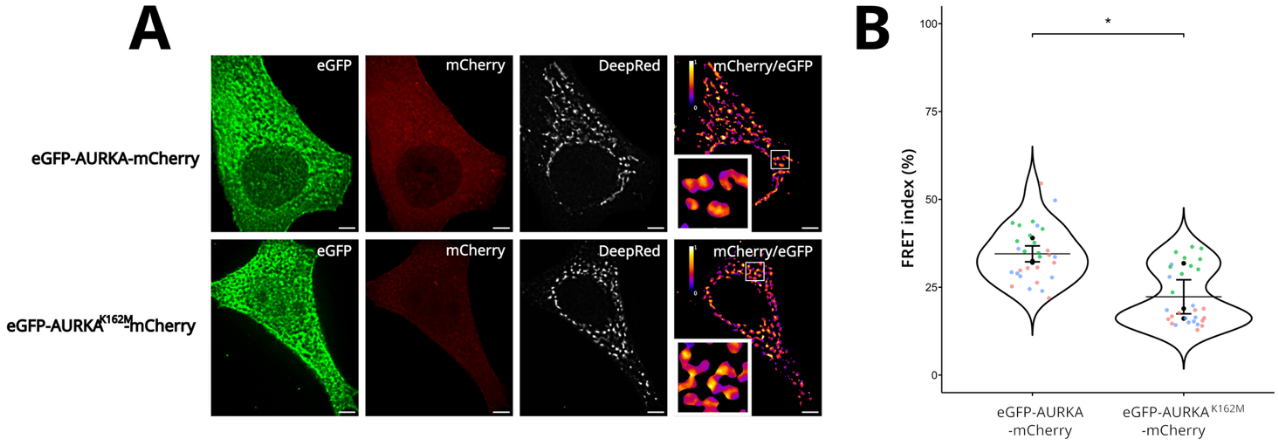
A
eGFP
mCherry
DeepRed
mCherry/eGFP
eGFP-AURKA-mCherry
eGFP-AURKA$^{K162M}$-mCherry
B
FRET index (%)
100
75
50
25
0
*
eGFP-AURKA
-mCherry
eGFP-AURKA$^{K162M}$
-mCherry

**Figure S4: The K162M mutation significantly lowers AURKA activation**

**(A)** Super-resolved SRRF images of the GFP, mCherry (FRET), Mitotracker DeepRed channels, and visual representation of ratiometric FRET in cells expressing AURKA or the K162M biosensor. **(B)** SuperPlots of FRET Index values (0-100 %) in cells expressing the AURKA biosensor or the K162M counterpart. $n$ = at least 10 cells per condition in each independent replicate. Pseudocolour scale on FRET images: ratiometric FRET (0-1). Data are means $\pm$ SEM. Each dot in SuperPlots represents FRET index values from individual cells sorted by replicate (colours). Tukey test: $^*P < 0.05$.

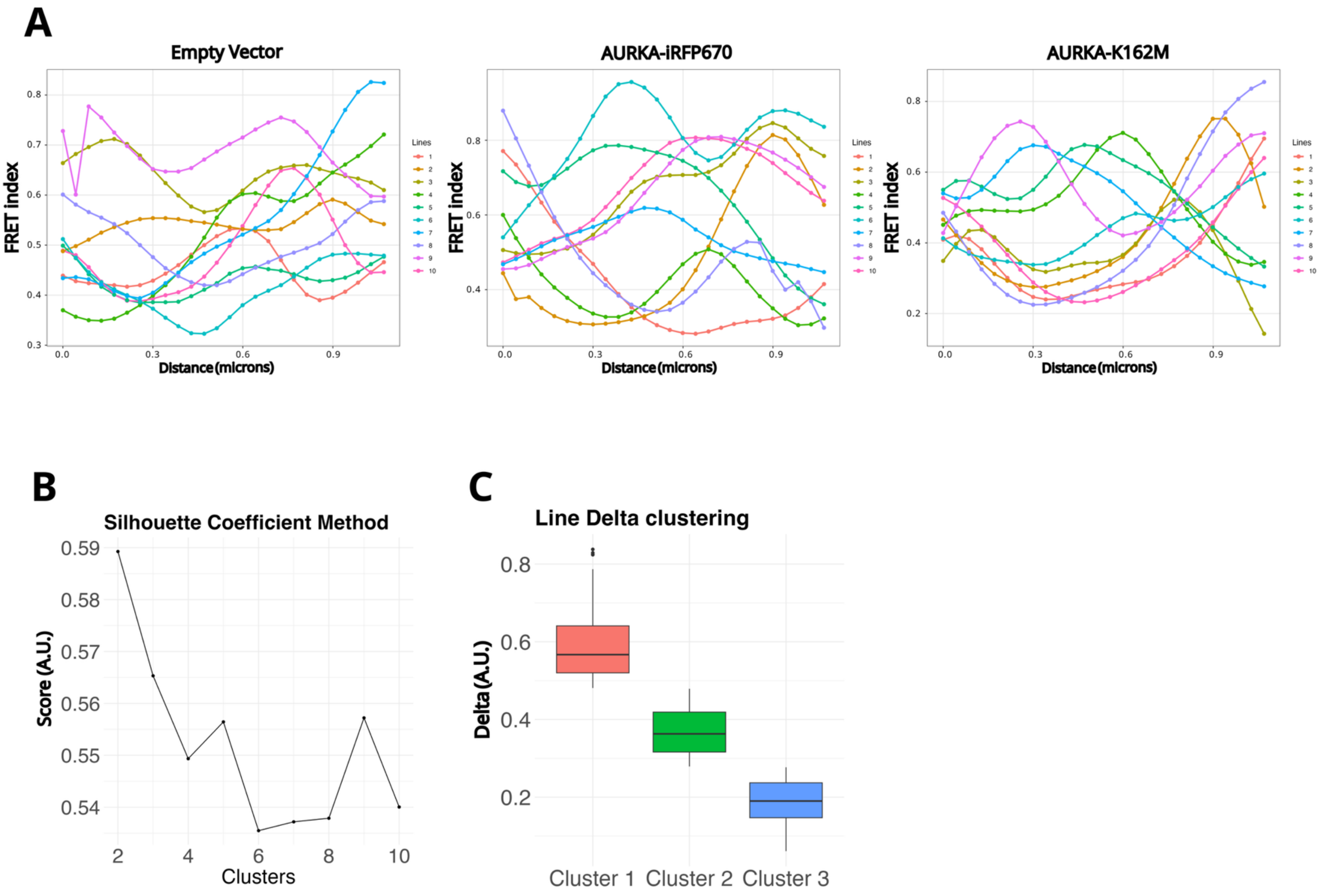
A
Empty Vector
AURKA-iRFP670
AURKA-K162M
FRET index
Distance (microns)
Lines
B
Silhouette Coefficient Method
Score (A.U.)
Clusters
C
Line Delta clustering
Delta (A.U.)
Cluster 1
Cluster 2
Cluster 3

**Figure S5: Random line analyses and clusters of MitoGO-ATeam2 co-expressed with AURKA or AURKA$^{K162M}$**

**(A)** Representative individual lines issued from the random line analyses. The graphs show FRET index values over distance (in µm) for each indicated condition. Each colourrepresents an individual random line. (**B**) Mean silhouette coefficient score (A.U.) as a function of the number of clusters. **(C)** Distribution of the delta of FRET index values across the three identified clusters. Data go from mean to max, and boxplots show the 1$^{st}$ and 3$^{rd}$ interquartile range and the median delta value contained within each boxplot. Clusters are ordered by decreasing median delta values.

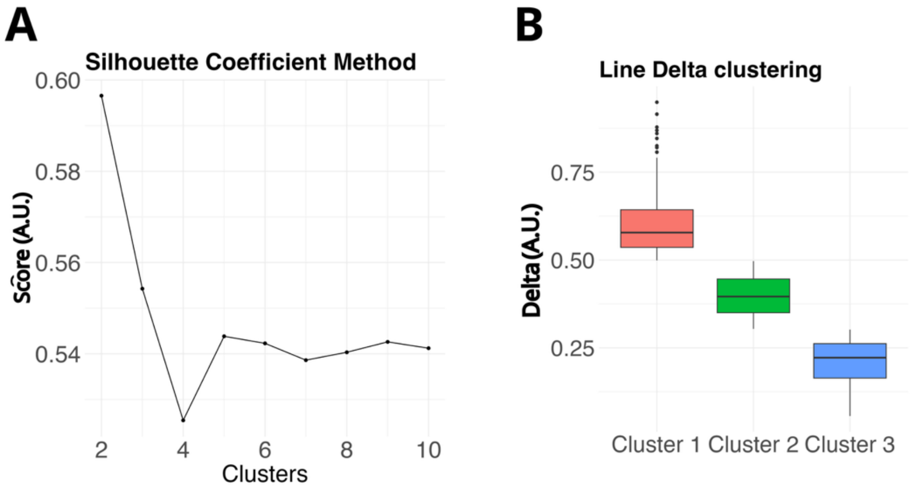
A
Silhouette Coefficient Method
Scôre (A.U.)
0.60
0.58
0.56
0.54
2
4
6
8
10
Clusters
B
Line Delta clustering
Delta (A.U.)
0.75
0.50
0.25
Cluster 1
Cluster 2
Cluster 3

**Figure S6: Random line analysis clusters of MitoGO-ATeam2 upon treatment with Xanthohumol**

**(A)** Mean silhouette coefficient score (A.U.) as a function of the number of clusters. **(B)** Distribution of the delta of FRET index values across the three identified clusters. Data go from mean to max, and boxplots show the 1$^{st}$ and 3$^{rd}$ interquartile range and the median delta value contained within each boxplot. Clusters are ordered by decreasing median delta values.

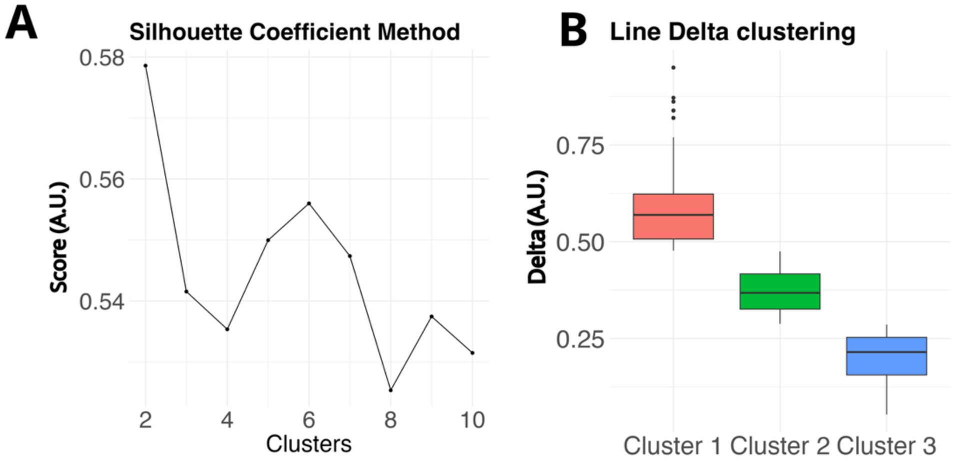
A
Silhouette Coefficient Method
0.58
0.56
0.54
Score (A.U.)
2
4
6
8
10
Clusters
B
Line Delta clustering
0.75
0.50
0.25
Delta (A.U.)
Cluster 1
Cluster 2
Cluster 3

**Figure S7: Random line analysis clusters of MitoGO-ATeam2 upon treatment with Compound 12**

**(A)** Mean silhouette coefficient score (A.U.) as a function of the number of clusters. **(B)** Distribution of the delta of FRET index values across the three identified clusters. Data go from mean to max, and boxplots show the 1st and 3rd interquartile range and the median delta value contained within each boxplot. Clusters are ordered by decreasing median delta values.

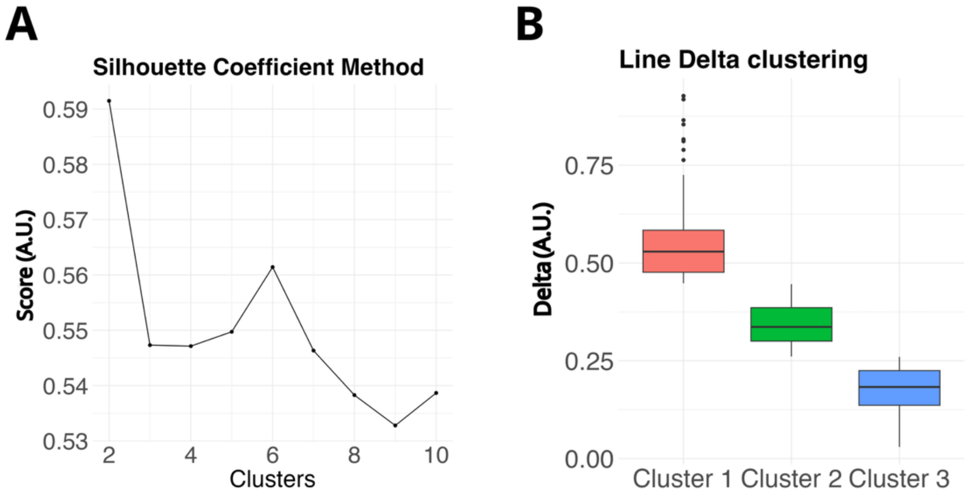
A
Silhouette Coefficient Method
Score (A.U.)
0.59
0.58
0.57
0.56
0.55
0.54
0.53
2
4
6
8
10
Clusters
B
Line Delta clustering
Delta (A.U.)
0.75
0.50
0.25
0.00
Cluster 1
Cluster 2
Cluster 3

**Figure S8: Random line analysis clusters of cells expressing the AURKA biosensor or its F31I counterpart**

**(A)** Mean silhouette coefficient score (A.U.) as a function of the number of clusters. **(B)** Distribution of the delta of FRET index values across the three identified clusters. Data go from mean to max, and boxplots show the 1$^{st}$ and 3$^{rd}$ interquartile range and the median delta value contained within each boxplot. Clusters are ordered by decreasing median delta values.

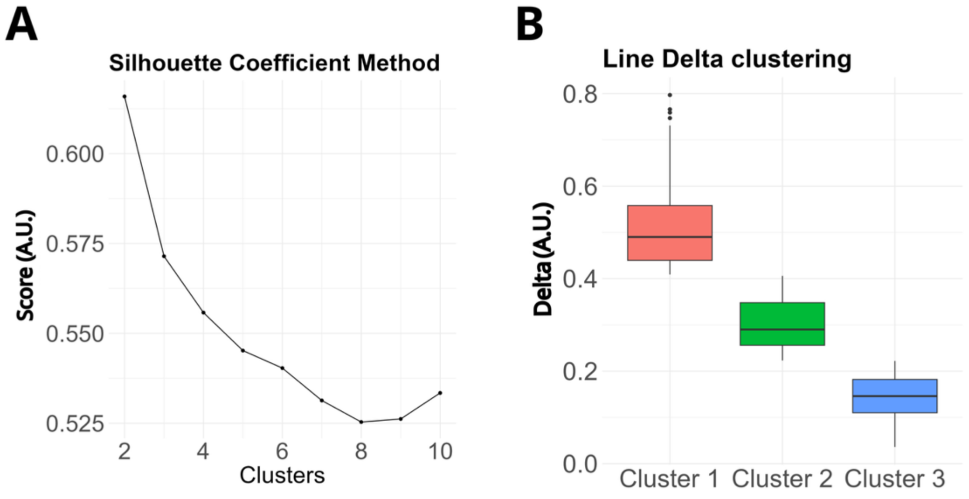

A
Silhouette Coefficient Method
Score (A.U.)
0.600
0.575
0.550
0.525
2
4
6
8
10
Clusters
B
Line Delta clustering
Delta (A.U.)
0.8
0.6
0.4
0.2
0.0
Cluster 1
Cluster 2
Cluster 3

**Figure S9: Random line analysis clusters of the MitoGO-ATeam2 in cells co-expressing AURKA or AURKA$^{F31I}$**

**(A)** Mean silhouette coefficient score (A.U.) as a function of the number of clusters. **(B)** Distribution of the delta of FRET index values across the three identified clusters. Data go from mean to max, and boxplots show the 1$^{st}$ and 3$^{rd}$ interquartile range and the median delta value contained within each boxplot. Clusters are ordered by decreasing median delta values.

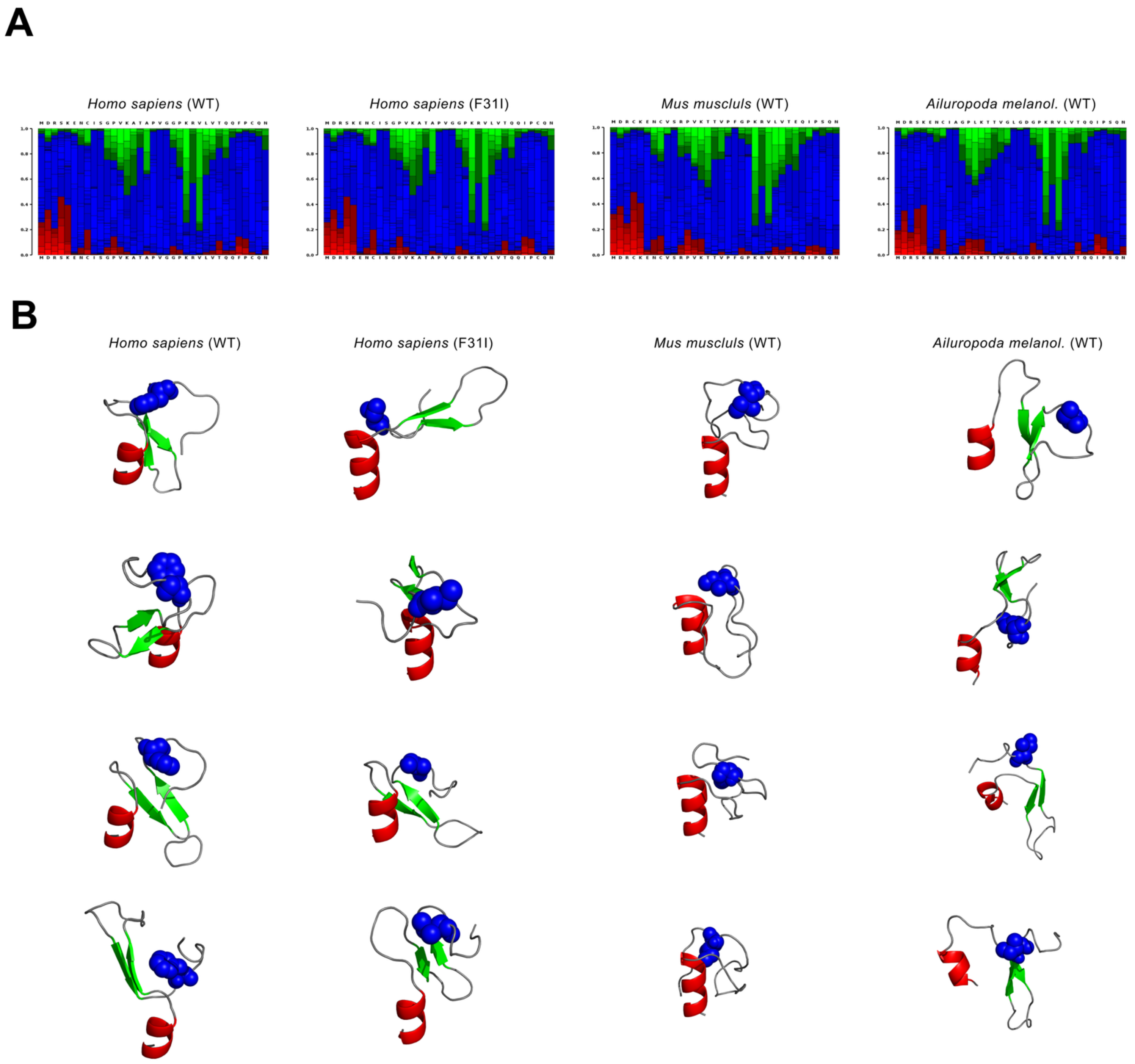
A
Homo sapiens (WT)
Homo sapiens (F31I)
Mus musculs (WT)
Ailuropoda melanol. (WT)
B
Homo sapiens (WT)
Homo sapiens (F31I)
Mus musculs (WT)
Ailuropoda melanol. (WT)

**Figure S10: PEP-Fold4 models of AURKA MTS**

**(A)** PEP-Fold4 confidence models on the structure motifs of the first model of the peptides generated. Colours: green: β-sheets, blue: coil, red: α-helix. **(B)** Representation of the next four 3D models generated by PEP-Fold4 for each MTS. Colours: green: β-sheets, grey: coil, red: α-helix, blue: Isoleucine or Phenylalanine residues.
